# A male-essential microRNA is key for avian sex chromosome dosage compensation

**DOI:** 10.1101/2024.03.06.581755

**Authors:** Amir Fallahshahroudi, Leticia Rodríguez-Montes, Sara Yousefi Taemeh, Nils Trost, Memo Tellez, Maeve Ballantyne, Alewo Idoko-Akoh, Lorna Taylor, Adrian Sherman, Enrico Sorato, Martin Johnsson, Margarida Cardoso Moreira, Mike J. McGrew, Henrik Kaessmann

**Affiliations:** Center for Molecular Biology (ZMBH), DKFZ-ZMBH Alliance, Heidelberg University, Heidelberg, Germany; Current affiliation: Department of Medical Biochemistry and Microbiology, Biomedical Center (BMC), Uppsala University, Uppsala, Sweden; The Roslin Institute and Royal (Dick) School of Veterinary Studies, University of Edinburgh, Edinburgh, UK; School of Biochemistry, Faculty of Life Sciences, University of Bristol, Bristol, UK; Reneco International Wildlife Consultants, Abu Dhabi, Abu Dhabi, United Arab Emirates; Department of Animal Biosciences, Swedish University of Agricultural Sciences, Uppsala, Sweden; Evolutionary Developmental Biology Laboratory, Francis Crick Institute, London, UK

**Author notes:** These authors jointly supervised this work (Mike J. McGrew, Henrik Kaessmann). **Corresponding authors** Correspondence to Amir Fallahshahroudi, Mike J. McGrew, or Henrik Kaessmann. e-mail: A.F.; M.J.M.; H.K.

## Abstract

**Birds have a sex chromosome system in which females are heterogametic (ZW) and males are homogametic (ZZ). The differentiation of avian sex chromosomes from ancestral autosomes entailed the loss of most genes from the W chromosome during evolution. However, to what extent mechanisms evolved that counterbalance the consequences of this extensive gene dosage reduction in female birds has remained unclear. Here we report functional in vivo and evolutionary analyses of a Z-chromosome-linked microRNA (miR-2954) with strongly male-biased expression that was previously proposed to play a key role in sex chromosome dosage compensation^1^. We knocked out miR-2954 in chicken, which resulted in early embryonic lethality of homozygous knockout males, likely due to the highly specific upregulation of dosage-sensitive Z-linked target genes of miR-2954. Our evolutionary gene expression analyses further revealed that these dosage-sensitive target genes have become upregulated on the single Z in female birds during evolution. Altogether, our work unveils a scenario where evolutionary pressures on females following W gene loss led to the evolution of transcriptional upregulation of dosage-sensitive genes on the Z not only in female but also in male birds. The resulting overabundance of transcripts in males resulting from the combined activity of two dosage-sensitive Z gene copies was in turn offset by the emergence of a highly targeted miR-2954-mediated transcript degradation mechanism during avian evolution. Our findings demonstrate that birds have evolved a unique sex chromosome dosage compensation system in which a microRNA has become essential for male survival.**

---

Sex chromosomes emerged from different sets of ancestral autosomes during the evolution of amniotes^2,3^ (Fig. 1a). These sex-chromosome formation events entailed substantial remodeling of gene contents and expression patterns due to structural changes and sex-related selective forces^3,4^. In particular, these events frequently involved massive losses of genes on the sex-specific chromosome; that is, the Y chromosome in male-heterogametic XY systems, and the W chromosome in ZW systems, where females are heterogametic^3,4^. As a consequence of the widespread gene dosage reductions in the heterogametic sex, compensatory mechanisms evolved^3-5^. For example, in the XY system of placental and marsupial (i.e., therian) mammals (Fig. 1a), dosage reductions in males were mainly compensated by an approximately twofold increase of expression levels for many of the genes on the single X chromosome through upregulation at both the transcriptional and translational layers^4-6^, which restored ancestral expression levels in males, but also other mechanisms^4,5^. The overexpression of X-linked genes in females from the combined activity of the two upregulated X chromosomes (i.e., the therian upregulation mechanisms are not specific to males) was then secondarily compensated by the process of X-chromosome inactivation, mediated by the *XIST* and *RSX* long noncoding RNAs (lncRNAs) in placentals and marsupials, respectively^7,8^. By contrast, in the XY system of green anole lizards (Fig. 1a), a male-specific mechanism of twofold transcriptional upregulation akin to that of fruit flies evolved^9^, rendering the evolution of female X inactivation unnecessary. Notably, in all of these systems, the combined actions of the different mechanisms have resulted in similar expression outputs in males and females for most X-linked genes^3,4^.

**Figure 1.**
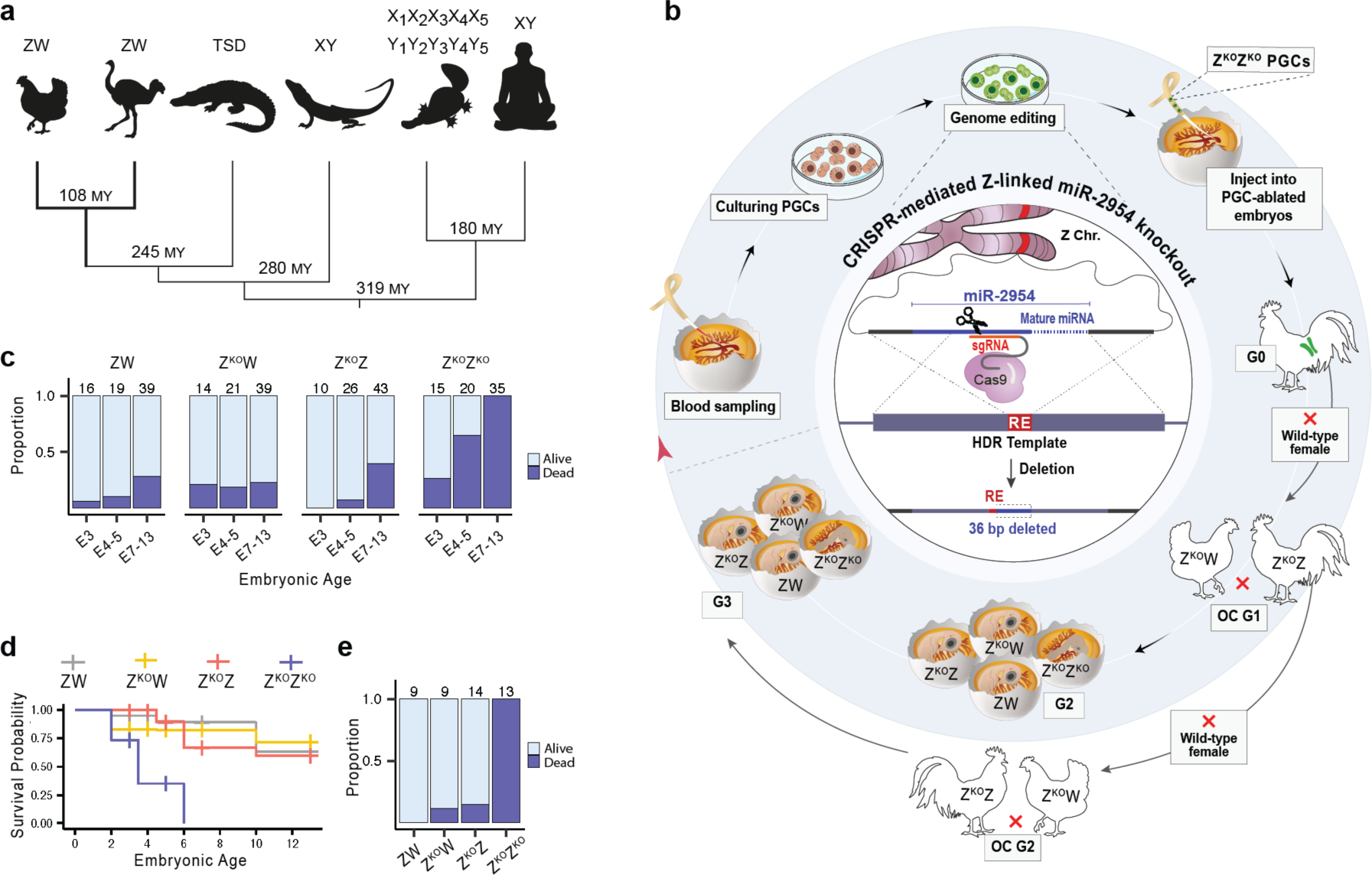
Role of miR-2954 in male chicken development. **a**, Overview of major known sex determination systems in amniotes^2,3,9^: ZW in birds (icons indicate chicken and ostrich, marking the deepest divergence in the bird phylogeny), temperature sex determination (TSD) in crocodiles, XY in Iguania lizards (icon reflects the green anole), multiple (5X and 5Y) sex chromosomes in platypus, and XY in humans. Approximate divergence times in millions of years (MY) are indicated at the respective nodes. **b**, Schematic of the experimental design used for the generation of miR-2954 knockout chickens across different generations (G0-G3), based on genome editing (CRISPR/Cas9 with single-guide RNA, sgRNA) in primordial germ cells (PGCs) and outcrossings (OC), and the assessment of resulting phenotypes (see Methods and Extended Data Fig. 1 for details). The restriction enzyme (RE) site used for genotype screening and the homology-directed repair (HDR) template are indicated. **c**, Distribution of alive and dead second-generation (G2) embryos, categorized by genotype (female wild-type ZW, female hemizygous knockout Z^KO^W, male heterozygous knockout Z^KO^Z, and male homozygous knockout Z^KO^Z^KO^) and embryonic day of development (E). Numbers above the bars represent the total count of embryos analyzed for each subgroup. **d**, Kaplan–Meier survival curve showing survival rates for embryos with different genotypes during development. **e**, Distribution of live and dead third-generation (G3) embryos at embryonic day 14, grouped by genotype.

In birds, however, which have a ZW sex chromosome system (Fig. 1a), the expression output of Z-linked genes is overall substantially higher in ZZ males compared to ZW females^5,10-12^, and the extent and importance of sex chromosome dosage compensation as well as the nature of potential underlying mechanisms have remained poorly understood. Previous work showed that Z-linked genes in female birds have only been partially and/or in part transcriptionally upregulated during evolution, rendering their transcript abundances lower than those of their evolutionary precursors on the ancestral autosomes^5,9^. Males have retained ancestral expression levels on their two Z chromosomes^5,9^, and so it has been unclear whether they have been affected by any upregulation mechanism. However, based on the observation that dosage sensitive genes show more similar expression levels between males and females, it was suggested that dosage compensation mechanisms in birds specifically target dosage sensitive genes^13^. Notably and consistently, we recently characterized a largely male-specific Z-chromosome-derived microRNA (miR-2954), whose predicted target genes, which were found to be mostly Z-linked, are enriched for dosage-sensitive genes and show significantly more similar transcript abundances between males and females than other genes on the Z chromosome^1^. Altogether, these observations suggested that miR-2954 might play an important role in avian dosage compensation^1^.

Here we assessed the function of miR-2954 *in vivo* using a chicken knockout model. In combination with evolutionary genomics analyses, this work enabled us to establish the mechanisms, extent and importance of avian dosage compensation.

## Generation of a miR-2954 chicken knockout line

To assess the functional role of miR-2954, which is a “mirtron“^14^ (i.e., it is located in an intron of the host gene *XPA*, Extended Data Fig. 1), we generated chicken knockout (KO) lines using genome editing of primordial germ cells^15-18^ (Fig. 1b) (Methods, Extended Data Fig. 1, and Supplementary Table 1). Briefly, we used a high-fidelity CRISPR-Cas9 system^19^, which minimizes potential off-target genomic alterations, and homology-directed repair^20^ to specifically knock out both copies of the miR-2954 locus (Z^KO^Z^KO^) in primordial germ cells (PGCs) derived from male embryos early in development^17,21^. We then generated a gonadal chimeric rooster (Generation 0; G0) by injecting these Z^KO^Z^KO^ PGCs into surrogate host chicken embryos whose own germ cell lineage was ablated in parallel^16,21^. One G0 rooster was raised to sexual maturity and mated with six wild-type hens, resulting in an outcross generation 1 (OC G1) with female hemizygous (Z^KO^W) and male heterozygous (Z^KO^Z) KO offspring. We observed no deleterious phenotypes in OC G1, and these birds reached sexual maturity and produced functional gametes, as evidenced by their ability to mate and produce viable offspring (Fig. 1b). Subsequently, we mated OC G1 males and females to produce second-generation (G2) embryos, resulting in the genotypes ZW, Z^KO^W, Z^KO^Z, and Z^KO^Z^KO^. Additionally, by mating an OC G1 male with six wild-type females, we generated a second outcross generation (OC G2). Offspring from the OC G2 were bred with each other to create third-generation (G3) embryos, affording confirmation of phenotypic observations made in the second generation (G2).

Our KO generation approach enabled us to investigate the phenotypic effects of the different genotypes in sibling birds, minimizing variability in the genetic background. Importantly, our repeated outcrossings also essentially rule out the presence of potential off-target mutations from the genome-editing procedure.

## miR-2954 is male-essential

To assess potential phenotypic consequences of the miR-2954 KO, we first assessed the viability of 297 G2 embryos on embryonic (E) days E3, E4-5, and E7-13 by assessing their morphology and heartbeat under a stereo microscope (Methods, Supplementary Table 2, and Supplementary Data 1). We found that wild-type female (ZW) chicken embryos as well as hemizygous female (Z^KO^W) and heterozygous male (Z^KO^Z) KO embryos exhibited similar average survival rates of approximately 79-85% (Fig. 1c, d, and Extended Data Fig. 2). These observations demonstrate that miR-2954 is not essential for female development, consistent with its very low expression in this sex^1^, and that the absence of one copy of the miR-2954 locus in males is not deleterious, implying that this microRNA is haplosufficient in male chicken embryos. In stark contrast to these genotypes, all males with the Z^KO^Z^KO^ genotype died before day 7 of embryonic development (Fig. 1c, d). When repeating these analyses for 45 embryos from day 14 of generation 3 (G3), we obtained very similar and hence confirmatory results, observing 100% lethality for Z^KO^Z^KO^ males and survival rates of around 86-100% for all other genotypes (Fig. 1e). We note that the fact that only homozygous KO embryos show a (lethal) phenotype, further rules out the presence/effect of potential off-target mutations from the genome-editing procedure, given that these would also affect heterozygous male or hemizygous female KOs.

Altogether, this work reveals that miR-2954 is male-essential and thus represents the only known sex-specific essential microRNA reported across taxa so far.

## Derepression of genes on the Z chromosome in KO animals

While microRNAs may also repress translation, they predominantly regulate the abundance of target messenger RNAs (mRNAs) by guiding Argonaute proteins to these targets through short (6-8 nucleotides, nt) seed sequences in the microRNA that are complementary to sites in the 3’ untranslated region (UTR) of the mRNAs, which are then degraded^14,22,23^. To understand the molecular basis of the male-lethal homozygous KO phenotype of miR-2954, we therefore sought to systematically assess transcript abundances of genes across the genome in KO chicken embryos compared to wild-type controls. To this end, we generated RNA sequencing (RNA-seq) data for both male and female chicken embryos as well as for the head, heart, and rest of the body of E3 and E5 males across all KO and wild-type genotypes, covering a total of 72 samples (Methods, Extended Data Fig. 3).

Based on these RNA-seq data, we first assessed changes in gene expression between samples from male homozygous (Z^KO^Z^KO^) KO and wild-type (ZZ) embryos. To assess the association of potential changes in expression with the loss of miR-2954 in the KO animals, we predicted its potential targets using TargetScan, which screens for the presence of short (6-8 nucleotide) UTR sequences that match a microRNA’s seed region^24^ (Methods, Supplementary Table 3). Our analyses revealed that predicted Z-linked and autosomal target genes of miR-2954 show significantly higher increases of transcript abundances in KOs compared to wild-type controls than non-target genes on the respective types of chromosomes (Fig. 2a, left and middle panels and Supplementary Table 4), in agreement with the gene repression role of microRNAs and the loss of repression in the miR-2954 KO embryos. Notably, however, the expression level increases in Z^KO^Z^KO^ samples of predicted Z-linked targets (median log_2_-fold change: whole embryo: 0.41, head: 0.45, heart: 0.48, body: 0.48) are substantially higher than those of predicted autosomal targets (median log_2_-fold change: whole embryo: 0.01, head: 0, heart: -0.01., body: 0.01), which are, overall, not or only slightly upregulated in the KO embryos compared to the wild-type controls (Fig. 2a, left panels).To identify the genes driving these patterns, we identified genes with significant expression differences between Z^KO^Z^KO^ and wild-type ZZ embryos using DESeq2 (ref. ^25^). We found that ∼50% of all 375 predicted Z-linked targets genes are significantly differentially expressed between the Z^KO^Z^KO^ and wild-type ZZ embryos (Fig. 2a, right panel) in the different tissues, whereas only ∼3-5% of the 3,383 predicted autosomal targets are differentially expressed, which, in turn, is significantly yet only marginally higher than the percentage observed for autosomal non-target genes (∼2-5%) (Fig. 2a). Moreover, the proportions of upregulated genes among differentially expressed predicted Z-linked target genes (∼99-100%) are significantly higher than those for differentially expressed predicted autosomal targets across tissues (∼38-54%) (all *P*-values < 10^-15^, two-sided ξ^2^ test). In further support of a key role of miR-2954, which is ubiquitously expressed in the body throughout development^1,26^, in repressing transcript abundances of Z-linked genes, we found that a large proportion (∼54%) of Z-linked target genes show consistent expression level increases in KO embryos across tissues – in stark contrast to predicted autosomal targets (Fig. 2b).

**Figure 2.**
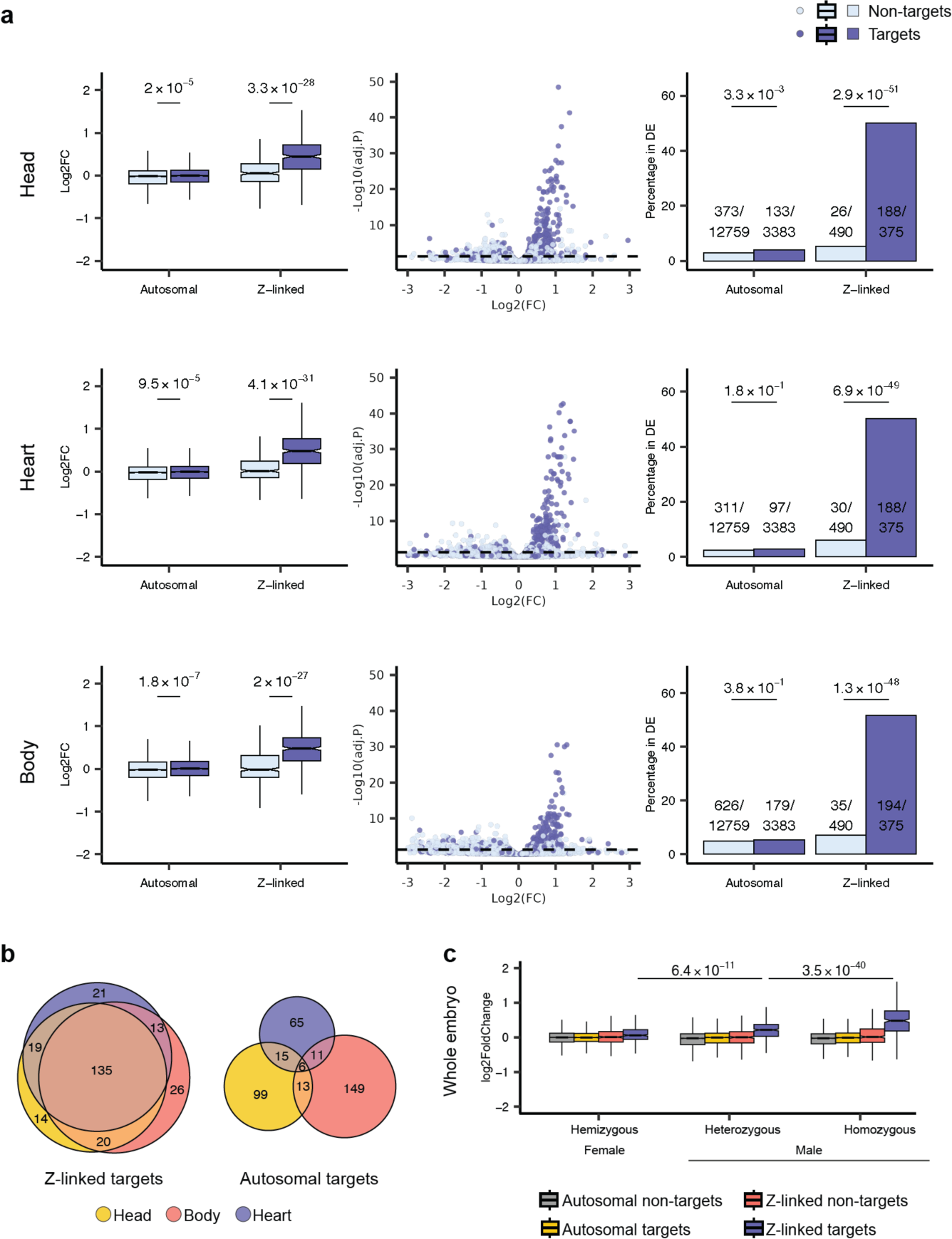
Impact of miR-2954 knockout on gene expression in different tissues. **a,** Left column: log_2_-fold changes (Log2FC) in gene expression between Z^KO^Z^KO^ and ZZ genotypes for both autosomal and Z-linked protein-coding genes; *P*-values from a two-sided Wilcoxon rank-sum test are indicated above. Log2FC estimates are based on transcriptomes of E3 and E5 embryos for the head, heart, and rest of the body, respectively. Middle column: volcano plots showing the log2FC and -log10 of Benjamini-Hochberg adjusted *P*-values for predicted miR-2954 target and non-target genes, contrasting Z^KO^Z^KO^ and ZZ genotypes. Right column: proportions of autosomal and Z-linked target and non-target genes among the differentially (DE) expressed genes (Benjamini-Hochberg adjusted *P* < 0.05) when contrasting Z^KO^Z^KO^ and ZZ genotypes. The distribution of predicted targets and non-targets of miR-2954 is compared using a chi-squared test for each group. **b**, Overview of the overlaps between Z-linked and autosomal DEGs, respectively, across different tissues. **c,** Log2FC in gene expression of autosomal and Z-linked target and non-target protein-coding genes for female hemizygous (Z^KO^W), male heterozygous (Z^KO^Z), and male homozygous (Z^KO^Z^KO^) genotypes compared to corresponding wild-type controls. *P*-values from a two-sided Wilcoxon signed-rank test represent the statistical significance of differences in Log2FC of Z-linked targets between Z^KO^W and Z^KO^Z, and between Z^KO^Z and Z^KO^Z^KO^ genotypes, respectively. Log2FC estimates are based on the transcriptome of whole embryos at E2.

We note that 26 to 35 predicted non-target Z-linked genes were also differentially expressed between the Z^KO^Z^KO^ and wild-type ZZ embryos, and that the proportions of upregulated genes among these genes (∼71-92%) are relatively similar to those of predicted Z targets (∼99-100%) and significantly higher than those of autosomal non-targets (∼38-54%) (all *P*-values < 10^-6^, two-sided χ^2^ test) (Fig. 2a). These observations are likely explained by false negative target predictions for a subset of Z-linked genes, due to, for example, non-canonical target sites (e.g., in the gene body), limited UTR annotations in the chicken genome and/or by repressive effects on the Z chromosome that occur downstream of the initial miR-2954-driven gene repression (e.g., upon the repression of transcription factors that regulate Z-linked effector genes). Thus, more Z-linked genes are probably regulated (directly or indirectly) by miR-2954 than predicted; interestingly, these may include *XPA*, the host gene of miR-2954, which is not a predicted target but shows a significant expression change (log_2_-fold change: ∼0.6-1.2) in KOs relative to controls (Supplementary Table 4).

Next, we investigated patterns of gene expression change in male heterozygote (Z^KO^Z) KOs. Akin to the pattern observed for homozygous (Z^KO^Z^KO^) KOs, this analysis revealed a predominant upregulation of predicted Z-linked target genes in Z^KO^Z animals (Fig. 2c; Extended Data Fig. 4). However, the extent of upregulation is significantly weaker in the heterozygote KOs (Fig. 2c), consistent with the presence of one remaining copy of the miR-2954 locus (i.e., partial repression of genes by this microRNA) and the lack of a (lethal) phenotype for Z^KO^Z embryos. Indeed, an analysis of small RNA-seq data that we generated for E5 males (Methods) confirms partial expression of miR-2954 in Z^KO^Z embryos, with levels that are 15-48% reduced compared to wild-type animals and, as expected, the complete absence of miR-2954 expression in Z^KO^Z^KO^ embryos (Fig. 3a). Notably, global analyses of the microRNA data confirm the high expression of miR-2954 in males^1^ and that other microRNAs are unaffected by its KO (Fig. 3b, Supplementary Table 5), further tying the expression and phenotypic effects in the KO animals directly to the loss of miR-2954-induced gene repression.

**Figure 3.**
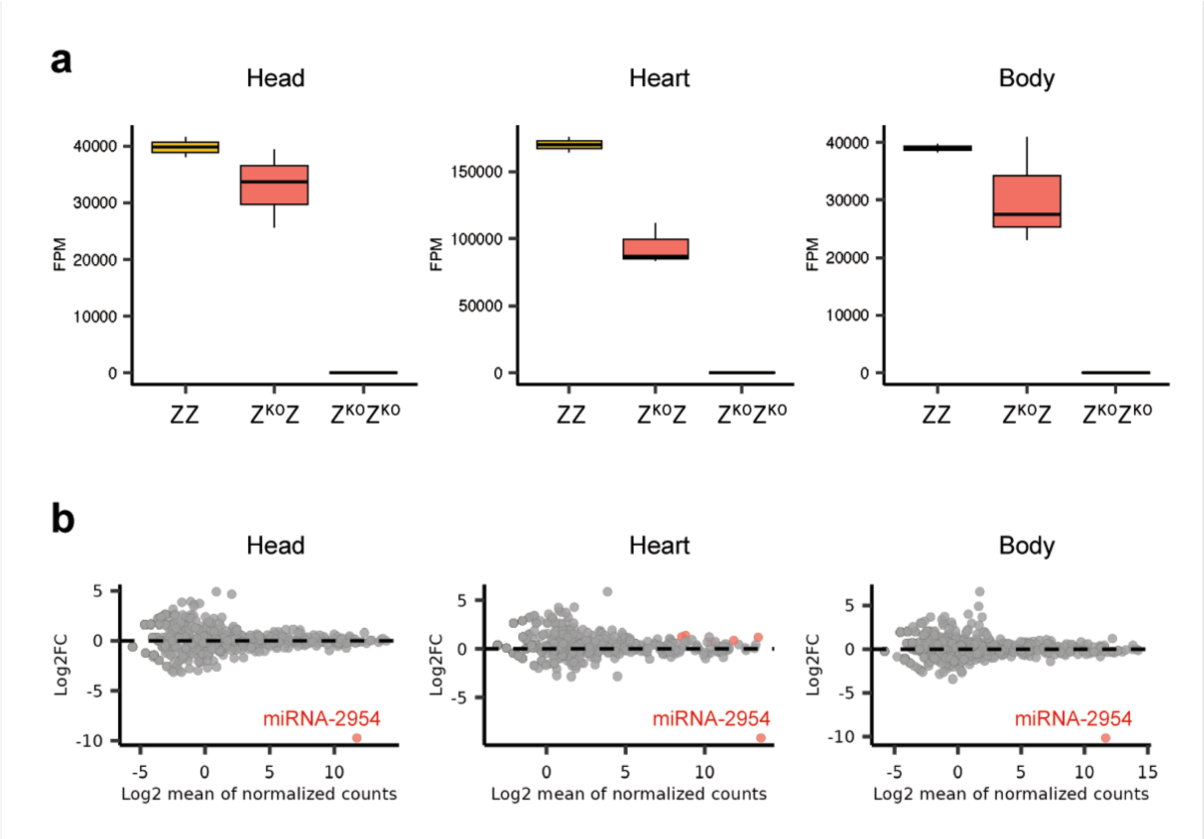
Impact of miR-2954 knockout on microRNA expression profiles in different tissues. **a**, Distributions of expression levels (fragments per million mapped reads, FPM) of miR-2954 for ZZ, Z^KO^Z, and Z^KO^Z^KO^ genotypes within different tissues at E5. **b**, MA plot visualizing mean expression and log_2_-fold change of miR-2954 and other mature microRNAs when comparing Z^KO^Z^KO^ and ZZ genotypes. MicroRNAs with significant expression changes are marked in red (Benjamini-Hochberg adjusted *P*-value < 0.01).

When investigating gene expression in female hemizygous (Z^KO^W) embryos, we again find a specific upregulation of Z-linked target genes compared to wild-type controls, but the pattern is even substantially weaker than that for male heterozygous (Z^KO^Z) embryos (Fig. 2c), in accord with the very low (i.e., on average 8-fold lower) expression of miR-2954 in females when compared to males^1^ and the absence of a deleterious phenotype in Z^KO^W female embryos.

We then sought to understand why miR-2954 preferentially targets Z-linked genes; that is, why a substantially higher proportion of predicted Z-linked targets compared to predicted autosomal targets become upregulated in KO animals. It is known that the potency of microRNA-mediated mRNA repression is greatly influenced by the degree of complementarity between the microRNA seed and the UTR of the target gene, with matching 8-mer sites being the most effective^14,24^. Furthermore, the presence of multiple binding sites for the same microRNA in a given target gene can amplify the repression effect^14,24^. Consistent with these modes of microRNA action, we find that predicted Z-linked targets are significantly enriched for 8-mer target sites and the presence of multiple target sites (of lengths between 6 to 8 nt) in their UTRs compared to predicted autosomal targets (Fig. 4a). In addition, we find that predicted Z-linked targets significantly upregulated in the KO animals are significantly enriched for 8-mer targets sites as well as multiple target sites relative to other predicted Z-linked targets (Fig. 4b).

**Figure 4.**
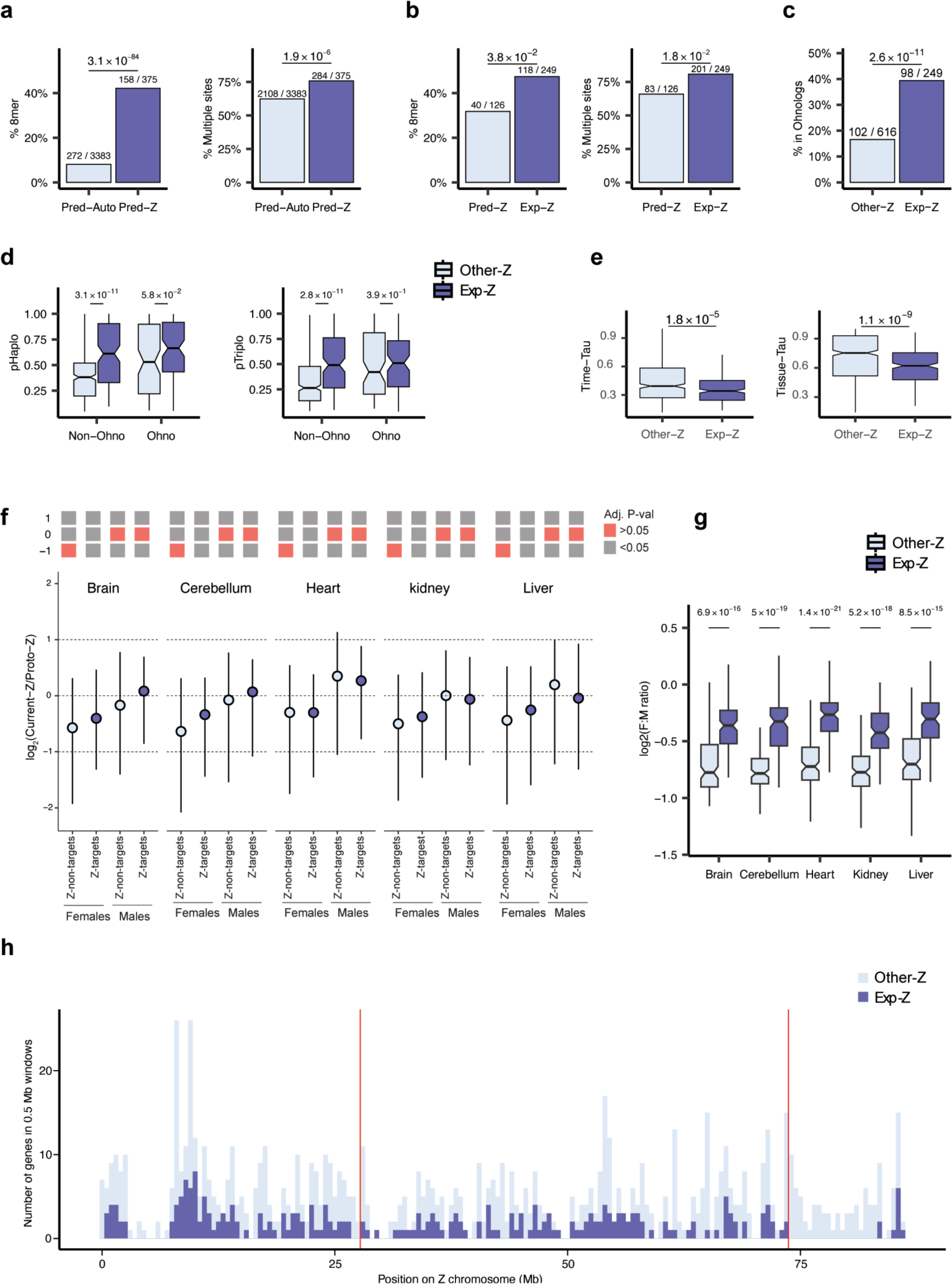
Experimental miR-2954 targets. **a**, Proportions of predicted autosomal (Pred-A) and Z-linked (Pred-Z) miR-2954 targets with 8-mer binding sites (left) and multiple binding sites (right); chi-squared *P*-values are indicated for the respective comparisons above the plots. **b**, Proportions of predicted Z-linked (Pred-Z) and experimentally validated Z-linked (Exp-Z) miR-2954 targets with 8-mer binding sites (left) and multiple binding sites (right; chi-squared *P*-values are indicated for the respective comparisons above the barplots. **c**, Proportions of experimentally validated Z-linked targets (Exp-Z) and other predicted Z-linked protein-coding genes (Other-Z) among chicken ohnologs; chi-squared *P*-values are indicated above the barplots. **d**, Probabilities of haploinsufficiency (pHaplo) and triplosensitivity (pTriplo) between Exp-Z and Other-Z genes for ohnologs and non-ohnologs, respectively; Wilcoxon test *P*-values are indicated above for the respective comparisons. **e**, Comparisons of tissue Tau scores and time Tau scores between Exp-Z and Other-Z genes, with Wilcoxon test *P*-values indicated above. **f**, Ratios of current versus proto-Z (ancestral) expression for Z-linked non-targets and Exp-Z genes for males and females, respectively. Values are plotted on a log2 scale to allow for linear and symmetrical patterns. Statistically significant deviations from the reference values (0.5 [log2 ratio of −1], current expression output is half of the ancestral one; 1 [log2 ratio of 0], current expression output is equal to the ancestral one; and 2 [log2 ratio of 1], current expression level is twofold higher than the ancestral one), as assessed by one-sample Wilcoxon signed rank tests (Benjamini-Hochberg corrected *P* < 0.05), are indicated above the plots (gray boxes; red boxes: no significant deviation). Error bars correspond to interquartile ranges. **g**, Female to male expression level ratios (log2) for Exp-Z and Other-Z genes, respectively, in different organs; Wilcoxon test *P*-values are indicated above for the respective comparisons. **h**, The number of Exp-Z and Other-Z genes in windows of 0.5 mega base pairs (Mb) along the Z chromosome; the position of male hypermethylated regions 1 and 2 (MHM1 and MHM2) are indicated by vertical lines.

Overall, the aforementioned results demonstrate that miR-2954 specifically targets Z-linked genes and that, therefore, its removal from the chicken genome leads to the upregulation of genes on the Z chromosome with lethal consequences in homozygous male KO embryos.

## miR-2954 targets dosage sensitive Z-linked genes

To understand why the upregulation of Z-linked targets of miR-2954 in Z^KO^Z^KO^ embryos is lethal, we set out to characterize these genes in detail. In these analyses, we focused on experimentally validated target genes, that is, predicted Z-linked target genes that show significant differential expression in at least one tissue of male homozygous KOs (Supplementary Table 3). The 249 targets (of which 248 are upregulated) account for ∼66% of all 375 predicted Z-linked targets and ∼29% of all 865 Z-linked protein-coding genes.

We explored whether experimental Z-linked targets are dosage sensitive, as would be expected in view of our observations that increases in transcript abundances for these genes have deleterious consequences in male homozygous KOs. Ohnologs^27^ are gene duplicates that are thought to typically be dosage sensitive, given that they have been retained as multiple paralogs after they emerged from the two rounds of whole-genome duplication in vertebrate ancestors (as first proposed by Susumu Ohno^28^) and are enriched for developmental functions and protein complex membership^29^. We find that experimental Z-linked targets encompass 49% (98/200) of all Z-linked ohnologs and that they are significantly enriched for this type of paralogs compared to other Z-linked genes (Fig. 4c).

However, not all dosage-sensitive genes are ohnologs and not all ohnologs may be dosage-sensitive. We therefore assessed dosage sensitivity across the entire Z chromosome using a resource that provides the probability of triplosensitivity (i.e., overexpression intolerance) and haploinsufficiency (i.e., deletion intolerance) for all human autosomal protein-coding genes^30^, which we transferred to the chicken genome via corresponding Z-linked 1:1 orthologs (Methods, Supplementary Table 3). Our analyses revealed that, as expected, chicken Z-linked ohnologs have significantly higher triplosensitivity and haploinsufficiency scores than other Z-linked genes (Fig. 4d). Importantly, these analyses also revealed that non-ohnolog Z-linked experimental targets also have significantly higher triplosensitivity and haploinsufficiency scores than other non-ohnolog Z-linked genes (Fig. 4d). This finding strongly supports a role for miR-2954 in regulating dosage-sensitive Z-linked genes in general – beyond ohnologs. Moreover, the pronounced upregulation of numerous dosage-sensitive genes on the Z chromosome likely explains the lethal phenotype observed in Z^KO^Z^KO^ embryos.

To further explore why the upregulation of Z-linked targets of miR-2954 in Z^KO^Z^KO^ embryos is lethal, we assessed their spatiotemporal gene expression patterns, given that the breadth of expression across tissues and developmental processes (here referred to as expression pleiotropy) is thought to represent a key determinant of the types of mutation that are permissible under natural selection^31^; that is, higher expression pleiotropy (broader expression) implies increased functional constraints. Indeed, when we assessed patterns of expression pleiotropy across Z-linked genes using a developmental transcriptome resource from our lab^32^ (Supplementary Table 6), we found that Z-linked targets have significantly broader spatiotemporal expression profiles than other Z-linked genes (Fig. 4e). This observation, together with the consistent repression of Z-linked targets by miR-2954, which is ubiquitously expressed across organs and development^1,26^ (Fig. 3b), provides a further explanation for the severe phenotype in the homozygous male KOs.

## Mechanisms of avian dosage compensation

The highly targeted and essential role of miR-2954 in repressing dosage sensitive Z-linked genes in males, together with the fact that Z-linked genes in females have only become partially or in part upregulated during evolution, whereas males retained ancestral expression levels^5,9^, are highly suggestive of an overall model for the evolution of highly targeted dosage compensation in birds. We hypothesized that the evolutionary pressures on females following W gene loss during the differentiation of avian sex chromosomes led to the evolution of transcriptional upregulation of dosage-sensitive genes on the single Z chromosome, which restored ancestral expression outputs (i.e., those on the ancestral autosomes) of these specific Z-linked genes in females. We further predicted that the underlying upregulation mechanism also affected males and that the resulting overabundance of transcripts in this sex resulting from the combined activity of two dosage-sensitive Z gene copies was, in turn, compensated by the emergence of the highly targeted miR-2954-mediated transcript degradation mechanism. In this scenario, the combined actions of both mechanisms (upregulation of dosage sensitive genes in females and their targeting by miR-2954 in males) should have resulted in more similar expression outputs for these dosage sensitive genes between the sexes (i.e., less reduced output in females) compared to other Z-linked genes.

To test this model of dosage compensation evolution, we first sought to assess whether dosage sensitive Z-linked genes that are targeted by miR-2954 (i.e., those that are upregulated in males upon KO of this microRNA) have become upregulated in females during evolution. To do so, we compared their current expression levels with their estimated ancestral levels prior to the formation of the sex chromosomes (using an RNA-seq dataset for adult organs^32^, Methods). We inferred ancestral expression levels of Z-linked genes from expression levels of autosomal 1:1 orthologs in an outgroup species (mouse) – an approach that has previously been shown to reliably approximate global patterns of ancestral expression^4,5,9,33^.

This analysis revealed that, in females, Z-linked miR-2954 target genes have indeed become significantly upregulated during evolution across different organs (i.e., current/ancestral expression output ratio > 0.5 [log_2_ ratio > −1]) – contrary to other Z-linked genes (Fig. 4f). In males, current expression levels are similar to ancestral levels (i.e., current/ancestral expression output ratio not significantly different from 1 [log_2_ ratio of 0]) for both Z-linked targets and other Z-linked genes, consistent with the maintenance of two Z copies for all genes in males and the repressive effect exerted by miR-2954 that compensates for the upregulation of the targets and is lost in our KO chicken model.

Next, we tested the last expectation of our model of dosage compensation evolution, that as a consequence of the different mechanisms of dosage compensation, the expression levels of Z-linked target genes should be more similar between females and males (i.e., less reduced in females compared to males) than for other Z-linked genes. Based on an extensive RNA-seq dataset for different organs across development^32^, we found that female-to-male expression level ratios are indeed significantly higher (i.e., female levels are closer to those of males) for Z-linked targets than for other Z-linked genes across all organs and developmental stages (Fig. 4g).

Finally, we note that when we explored the localization of Z-linked targets on the Z chromosome, we found that they are relatively evenly distributed along the Z chromosome (Fig. 4h), that is, without any overt clustering around the two small previously reported male hypermethylated (MHM) regions specific to galliform birds (i.e., chicken and turkey) that have been proposed to represent specific dosage compensated regions in this avian lineage^34,35^.

Altogether, our analyses provide strong support for our model for the evolution of dosage compensation of the ZW sex chromosome system, where W gene loss in females drove the evolution of transcriptional upregulation specifically of dosage sensitive Z-chromosomal genes along the Z chromosome in both sexes as well as the evolution of a secondary mechanism that mitigates transcript overabundances in males through miR-2954 (Fig 5a).

**Figure 5.**
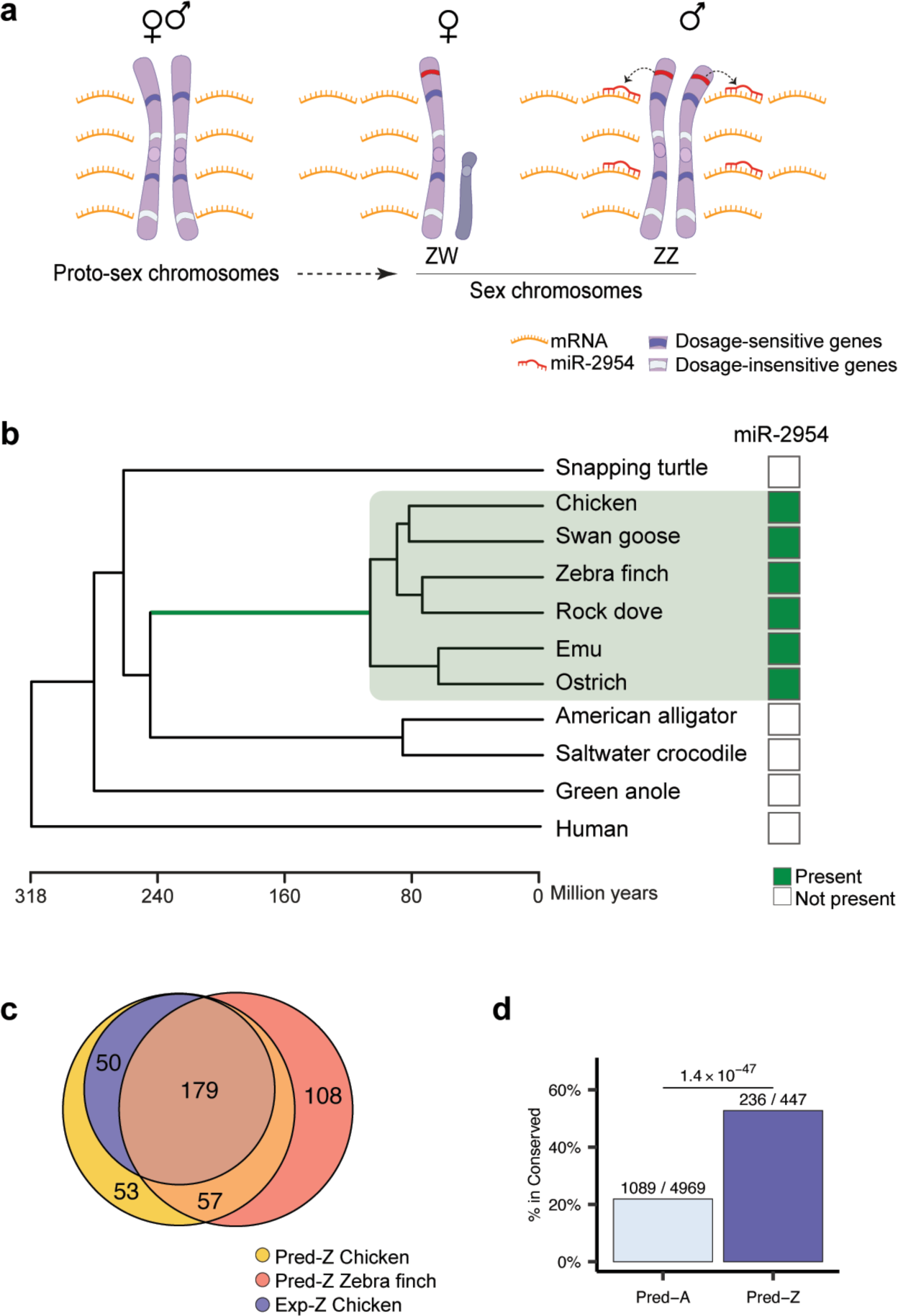
Evolution of avian dosage compensation and the emergence and conservation of miR-2954 and its targets. **a**, Illustration of the upregulation of dosage-sensitive Z-linked genes in birds following W gene loss and the concurrent evolution of miR-2954 to mitigate excess transcript accumulation in males. **b**, Phylogenetic distribution of miR-2954; the phylogenetic tree includes representative avian and other amniote species (branch lengths scaled to represent evolutionary time in millions of years). miR-2954 originated in the avian stem lineage (green branch). **c**, Overview of miR-2954 targets that are shared between experimentally validated targets in chicken (Exp-Z) and predicted targets in chicken and/or zebra finch (Pred-Z). **d**, Proportions of miR-2954 target genes on autosomes (Pred-A) and Z chromosome (Pred-Z) that are shared between chicken and zebra finch; the chi-squared test *P*-value for the comparison is indicated above.

## Conservation of miR-2954-mediated dosage compensation across birds

We finally sought to investigate the origin and evolution of this avian dosage compensation system. To this end, we first screened for the presence of the miR-2954 locus across amniotes (Methods). Notably, miR-2954 is present across all bird genomes but not in the genomes of any other species (including crocodiles – the closest living bird relatives), which, however, all contain the host gene of miR-2954 in birds, *XPA* (Fig. 5b). Strikingly, not only the mature sequence (22 nt) of miR-2954 but also its precursor sequence (68 nt) show 100% sequence conservation across all birds (Extended Data Fig. 5). Together, these observations suggest that the dosage compensation system, including the secondary compensatory mechanism mediated by miR-2954, emerged in the avian stem lineage between approximately ∼108-245 million years ago^36^ and has since been strongly selectively preserved. This finding is notable in view of the extensive diversity among birds in the degree of ZW differentiation^37^. That is, it suggests that the evolutionary pressures arising from sex chromosomal dosage imbalances have been strong in all bird lineages/species, even in those with less pronounced sex chromosome differentiation^37^, such as the flightless ratites (i.e., Paleognathae, such as emu and ostrich)(Fig. 1a and 5b).

To further assess the conservation of miR-2954-mediated dosage compensation across birds, we compared aspects of this mechanism between chicken and zebra finch, which represent two major avian lineages (Galliformes/Anseriformes and Neoaves, respectively) that diverged ∼91 million years ago^36^ and mark the deepest split within Neognathae. First, we found that ∼78% of the experimental Z targets we identified in chicken are also computationally predicted targets of miR-2954 in zebra finch, a species for which strongly male-biased expression of miR-2954 was previously shown as well^38^ (Fig. 5c). Moreover, in zebra finch, Z-linked computationally predicted targets are significantly more likely to also be targets in chicken than autosomal predicted targets (Fig. 5d). Together with our previous finding that predicted Z-linked targets (rather than autosomal predicted targets) and in particular Z-linked ohnologs are predominantly upregulated upon miR-2954 knockdown in a male zebra finch cell line^1^, these observations strongly support that the functional role of miR-2954 in zebra finch is similar to that in chicken; they thus also support the notion that the dosage compensation system is similar and indeed highly conserved across birds.

## Discussion

In this study, we have unraveled the function of miR-2954, based on a chicken KO model, as well as the evolutionary events and mechanisms of avian dosage compensation associated with this microRNA’s emergence. We unveiled a scenario where, driven by the decay of the W chromosome, dosage sensitive genes became upregulated on the Z chromosome not only in (ZW) females but, as a secondary consequence, also in (ZZ) males. The resulting overabundance of Z transcripts from these genes in males was, in turn, compensated by the evolution of a highly targeted miR-2954-mediated transcript degradation mechanism. During the co-evolution of miR-2954 and its targets, high and broad expression levels of miR-2954 emerged in males in concert with potent and, frequently, multiple target sites in the dosage sensitive genes it regulates, leading to the effective and substantial suppression of the target genes across the body and hence bringing their transcript levels back to their ancestral levels.

Our findings suggest that the miR-2954 mechanism has been highly conserved across birds, consistent with its crucial role in avian males. Indeed, our chicken experiments demonstrate that miR-2954 is essential for male viability, with the complete KO of miR-2954 leading to early embryonic lethality because of the upregulation of the large number of dosage sensitive genes it normally suppresses. The observation of a sex-specific essential role for a microRNA is remarkable, not least because KOs of single microRNAs have rarely had strong or even lethal phenotypes; that is, redundancy among microRNA family members often requires disruption of multiple family members before phenotypes become apparent^14^.

The avian ZW dosage compensation system we have uncovered here has key parallels to the XY system of therian mammals^3,4,6^ and Ohno’s original reasoning^39^. In both systems, gene expression upregulation was initially driven by the decay of the sex chromosome specific to the heterogametic sex – the W in female birds and the Y chromosome in male mammals. The fact that – unlike in the XY systems of the anole lizard^9^ or fruitflies^40^ – these upregulation mechanisms were not specific to the heterogametic sex, necessitated the evolution of secondary silencing mechanisms in male birds and female mammals^3^, which, however, show striking differences, even though both involve noncoding RNAs as key mediators and are essential for the viability of the heterogametic sex. While birds evolved a highly targeted and efficient post-transcriptional mechanism that specifically silences upregulated dosage sensitive genes on the Z chromosome, the more “radical” transcriptional silencing mechanisms in therian mammals, mediated by the *XIST* and *RSX* long noncoding RNAs in placental mammals and marsupials, respectively^3,8,41^, involve a transcriptional shutdown of most of the X chromosome. Future work may explore the reasons for these different evolutionary solutions to the secondary sex chromosome dosage imbalances in the homogametic sexes of birds and therian mammals, which might, for example, be related to differences in the proportions of dosage sensitive genes on the different ancestral autosomes.

Altogether, our work has identified a crucial and hitherto undescribed functional role for a microRNA in dosage compensation. Thus, rather than acting as a “sculptor” of the transcriptome^14^, like many microRNAs that have important but more subtle regulatory effects on gene expression^14^, miR-2954 has evolved as the key mediator of a targeted post-transcriptional silencing network that has ensured the survival of males in the wake of avian sex chromosome differentiation.

## Acknowledgement

We thank all members of the Kaessmann group for discussions; D. Ibberson and B. Berki from the Deep Sequencing Core Facility of Heidelberg University for sequencing support; D. Meunier, K. Hogan, F. Thomson, N. Russell, and the Roslin poultry team in the National Avian Research Facility for supporting the chicken in vivo work; and M. Clinton, M. Davey, and L. McTeir for useful discussions and support. This work was supported by grants from the European Research Council (ERC) to H.K. (grant agreement numbers 101019268 and 615253), by grants from the Swedish Research Council (VR) (2017-06218) and the Swedish Research Council for Sustainable Development (Formas) (2021-00513) to A.F., by a Formas grant (2023-01396) to S.Y.T., and by an add-on fellowship of the Joachim Herz Stiftung to L.R.-M. M.C.-M. was supported by the Francis Crick Institute, which receives its core funding from Cancer Research UK (grant CC2185 to M.C.-M.), the UK Medical Research Council (grant CC2185 to M.C.-M.), and the Wellcome Trust (grant CC2185 to M.C.-M.). This work was also supported by the Institute Strategic Grant Funding from the BBSRC (BB/P0.13732/1 and BB/P013759/1) to the Roslin Institute.

## Author contributions

A.F., M.J.M., and H.K. conceived and organized the study. A.F. and H.K. wrote the manuscript, with input from all authors. A.F. performed the majority of the experiments and bioinformatics analyses. L.R.-M. performed evolutionary and spatiotemporal expression analyses. E.S. performed the embryo survival analysis. S.Y.T. supported the experimental work. M.T.J., M.B., A.I.A., L.T., and A.S. made contributions to the experimental procedures. N.T., E.S., M.J., and M.C.-M. provided key feedback and discussions.

## Data availability

Raw mRNA and short RNA sequencing data have been deposited in SRA, NCBI (BioProject: PRJNA1079296) Reviewer link: https://dataview.ncbi.nlm.nih.gov/object/PRJNA1079296?reviewer=kcmuga1eo2up43sdj1t53shu5 Processed mRNA and short RNA sequencing data have been submitted to the Gene Expression Omnibus (GEO) repository and are currently processed. In the meantime, the processed data can be accessed here: https://drive.google.com/drive/folders/1XvM4o1M-hPcFNaPbkGAVTiyiD4_tToX-?usp=sharing

## Code availability

Custom scripts used to generate the results reported in the paper and processed data are available at github (https://github.com/amirshahr/MIR2954).

## Methods

### Isolating, sexing, and culturing PGCs

Genome editing in chickens involves the derivation and culturing of primordial germ cells (PGCs), performing genome editing on these cells, and the subsequent injection of the edited cells into surrogate hosts depleted of their native PGCs^21^. Following the injection of the genetically edited PGCs into the gonads of sterile surrogate hosts, the resulting offspring will inherit the genetic modifications introduced into the PGCs^21^ Fig. (1b and Extended Data Fig. 1). To establish miR-2954 KO lines, ten PGC lines were derived from the blood of Hy-Line Brown (HL) chicken embryos at Hamburger-Hamilton (HH) stage 16 (E2.5) and cultured according to previously described methods^21^. The sex of the PGC lines was determined according to previous work^21,42^ based on two sets of primers for one W-chromosome-specific gene and one autosomal gene (the control), respectively; the latter serves as a control for PCR reaction success (Supplementary Table 1). We cultured four male PGC lines and subsequently randomly selected one line for the knockout experiment. This PGC line was cultured for 22 days in total before transfection.

### Design of single-guide RNA (sgRNA) and homology-directed repair template

Inducing double-stranded breaks (DSBs) at specific genomic loci, followed by homology-directed repair using a template, introduces precise nucleotide substitutions^20^. Using CHOPCHOP v2 (ref. ^43^), we designed and tested five custom sgRNAs (Supplementary Table 1) to target the miR-2954 (MIR2954) locus (Gene ID: 100498678), located within the second intron of the DNA damage recognition and repair factor gene, *XPA* (ENSGALG00010009534), on the forward strand of chromosome Z (Location: NC_052572.1: 71305174-71305241; reference genome: bGalGal1.mat.broiler.GRCg7b (GCF_016699485.2). Additionally, we designed one single-stranded DNA oligonucleotide (ssODN) sequence as a repair template to exploit the homology-directed repair pathway. The ssODN repair template consisted of Ultramer DNA Oligonucleotides, custom-synthesized by Integrated DNA Technologies. The ssODN template contained homology arms flanking miR-2954, designed specifically for introducing a 36 bp deletion encompassing the entire mature miR-2954 sequence and part of its flanking pre-microRNA sequence. Additionally, we incorporated an EcoRI restriction endonuclease site (5’-GAATTC-3’) into this ssODN (Supplementary Table 1). These modifications effectively knock out miR-2954 and allow PCR-based genotyping for successful deletion events in both PGCs and the derived chickens (Extended Data Fig. 1 and Supplementary Table 1).

### Genotyping

We designed PCR primers to amplify a 550 bp region within the second intron of the *XPA* gene, encompassing the targeted deletion site (Supplementary Table 1) using Primer-Blast^44^. EcoRI restriction endonuclease enzyme specifically recognizes and cuts DNA at the restriction site (5’-GAATTC-3’). Following EcoRI digestion of this PCR product and subsequent gel electrophoresis, we expected to observe a single 550 bp band in wildtype individuals (ZZ and ZW) due to the absence of the EcoRI restriction site, three bands (550 bp, 298 bp, and 221 bp) in heterozygote knockout individuals (Z^KO^Z) due to digestion of half of the product, and two bands (298 bp and 221 bp) in homozygote males (Z^KO^Z^KO^) and hemizygote females (Z^KO^W) due to complete EcoRI restriction site digestion. This differential PCR band pattern served as a molecular signature for genotyping the individuals. The PCR was performed using Phusion High-Fidelity PCR Master Mix with GC Buffer from New England Biolabs, in accordance with the manufacturer’s guidelines. The reaction mixture was prepared with 1.25 µl of 10 µM Forward Primer, 1.25 µl of 10 µM Reverse Primer, 0.75 µl DMSO, 12.5 µl of 2X Phusion Master Mix, and ∼100 ng of DNA in 1 µl of water. The thermal cycling conditions were set as follows: an initial denaturation at 98°C for 60 seconds, followed by 35 cycles of 98°C for 10 seconds, 62°C for 20 seconds, and 72°C for 20 seconds, concluding with a final extension at 72°C for 10 minutes. To perform genotyping, we first extracted DNA from approximately 10,000 PGCs or embryonic tissues using DNeasy Blood & Tissue Kits from Qiagen, according to the manufacturer’s protocol. We then conducted PCR reactions as described above and subjected the PCR products to EcoRI digestion using EcoRI-HF and rCutSmart buffer from New England Biolabs, following the manufacturer’s guidelines. Each reaction consisted of 5 µl of PCR product, 1 µl of EcoRI-HF, 1 µl of rCutSmart buffer, and 8 µl of water, and was incubated at 37°C for 30 minutes, followed by a 5-minute heat inactivation at 65°C. Alternatively, the genotypes of multiple samples were analyzed based on the size of the undigested PCR products using the Agilent Fragment Analyzer system. In this approach, a 550 bp band represented ZZ and ZW, a 520 bp band represented Z^KO^Z^KO^ and Z^KO^W, and two bands (550 bp and 520 bp) in Z^KO^Z.

### PGC transfection, selection, and clonal expansion

We used a High-fidelity Cas9 variant (SpCas9-HF1), which significantly reduces off-target effects compared to wildtype Cas9^19^. For the expression of SpCas9-HF1 and sgRNAs in PGCs, we utilized the HF-PX459 (V2) expression vector, which also bears puromycin resistance as an antibiotic selection gene^17^ (Addgene plasmid #118632). We cloned all five sgRNAs individually into the plasmids according to previous descriptions^17,20^ and then tested the effectiveness of three of these plasmids harboring sgRNAs 1-3. We transfected 1.5 µg of the vector and 0.5 µg of ssODNs into approximately 100,000 HL PGCs using Lipofectamine 2000 transfection reagent (Thermo Fisher Scientific). After 24 hours in culture, the cells were treated with 0.6 µg/ml puromycin for 48 hours for the selection of successfully transfected cells. We then cultured these cells for around two weeks and then genotyped them for the presence of knockout PGCs via EcoRI digestion of the PCR product. Using gRNA3, we observed a strong PCR band at 550bp and two faint bands at approximately 300bp and 220bp. This pattern suggested the incorporation of the ssODNs template in a subset of transfected PGCs. Accordingly, these PGCs were sorted using the BD FACSAria III Cell Sorter (BD Biosciences) into a 96-well plate at a rate of one cell per well to identify the clonal populations with the deletion of miR-2954. After three weeks of culturing, we screened the genotypes of 42 clonal PGC populations that survived and propagated, and identified four Z^KO^Z and two Z^KO^Z^KO^ clonal populations among them (i.e., 6/42 clones were targeted). Subsequently, we cryopreserved the homozygote and heterozygote populations following established protocols^21^ and used one of the Z^KO^Z^KO^ populations for confirmation of the deletion and injection to surrogate hosts to generate the knockout animals. To confirm the deletion of miR-2954, we performed a PCR reaction on the DNA obtained from the PGC line before transfection. The selected clonal Z^KO^Z^KO^ PGC population and the resulting PCR products were sequenced by Eurofins Genomics (Ebersberg, Germany), using their Sanger sequencing services (TubeSeq Service). Analysis of the sequences confirmed the deletion of miR-2954 and integration of the EcoRI site as per the design of the provided ssODNs repair template (Extended Data Fig. 1).

### Generation of G0 rooster by injection of the PGCs into a surrogate host

The injections of the Z^KO^Z^KO^ PGCs into surrogate host embryos were done using our previously described method^21^. Briefly, we thawed the cryopreserved clonal Z^KO^Z^KO^ PGCs seven days prior to the intended injection date and propagated them to a density of approximately 150,000 cells per well in a 24-well tissue culture plate. These cultured PGCs were pelleted by standard centrifugation and then resuspended in the PGC culture medium to achieve a concentration of 5000 cells/μl. To this suspension, we added 0.1 μl of the chemical compound AP20187 (B/B) (25 mM) per 5 μl of PGC suspension. Approximately 1 μl of this mixture was aspirated into a microcapillary injection tube and injected into each iCaspase9 sterile embryo^16^ at HH stages 15–16. AP20187 (B/B), present in the injected PGC mixture, induces the dimerization of the FK506-binding protein (FKBP), leading to the activation of the attached caspase-9 protein and the induced apoptotic cell death of the native PGCs in the iCaspase9 sterile embryos, thereby allowing colonization of the gonads by the injected Z^KO^Z^KO^ PGCs^16^. Injecting the clonal PGCs into 20 iCaspase9 sterile embryos resulted in hatching of 7 G0 chicks, comprising 1 male and 6 females.

### Generation of miR-2954 knockout chickens

We maintained the male G0 and raised it to sexual maturity. This G0 was then paired with six HL hens (same breed), producing Z^KO^Z and Z^KO^W individuals (outcross generation 1; OC G1). We then raised five male and six female OC G1 individuals to sexual maturity. One of these males was mated with the OC G1 females to generate second-generation (G2) embryos (Z^KO^Z, Z^KO^Z^KO^, ZW, or Z^KO^W), used for viability studies and tissue collection for gene expression analyses. A second OC G1 male, not involved in generating G2 individuals, was mated with six HL females. This pairing produced OC G2 individuals for the genotypes ZZ, Z^KO^Z, ZW, and Z^KO^W. Finally, upon reaching sexual maturity, a OC G2 Z^KO^Z rooster was mated with six OC G2 Z^KO^W hens to produce G3 embryos (Z^KO^Z, Z^KO^Z^KO^, ZW or Z^KO^W). These G3 embryos were then used to confirm the phenotypes observed in the G2 generation (Fig 1a). All animal management, maintenance, and embryo manipulation were conducted under UK Home Office license PP9565661 and in accordance with all regulations. The experimental protocols and studies received approval from the Roslin Institute Animal Welfare and Ethical Review Board Committee.

### Measuring viability and sample collection

We assessed the viability of 297 G2 embryos and 45 G3 embryos across various developmental stages. For the G2 embryos, evaluations were conducted at embryonic (E) days E3-5, E7, and E13. At E3-5, we assessed normal development and the presence of a heartbeat using a stereo microscope. For stages E7-13, we assessed normal development and the presence of blood flow after removal of the chorioallantoic membrane. We collected E2 embryos (G2) for gene expression analysis but did not need to perform survival analysis at this stage to confirm the viability given the lack of heart development in these embryos. We evaluated the viability of G3 embryos at E14 following observation of blood flow after removal of the chorioallantoic membrane; however, a subset of the evaluated G3 embryos was frozen for subsequent phenotypic analyses based on their developmental progress. For all assessed embryos (dead and alive), we further captured stereo microscope images (SI) and collected samples of the extraembryonic membranes and vasculatures for molecular sexing and genotyping (as described above). We used these images to confirm our initial phenotypic assessment (Supplementary Table 2 and Supplementary Data 1).

Additionally, given that the G2 embryos were of the Z^KO^Z, Z^KO^Z^KO^, ZW, or Z^KO^W genotypes and did not include wild-type homozygous males (Supplementary Table 2), we added wild-type eggs to each experimental batch to include ZZ male embryos in the gene expression analyses. These HL eggs were derived from the same HL breed used to generate the G2 embryos, ensuring genetic consistency between the groups. All collected living G2 embryos were frozen on dry ice and subsequently preserved at -80°C for future dissections and RNA extraction in preparation for RNA sequencing.

### Survival analysis

We analyzed the survival of G2 embryos using proportional hazards models in the R package icenReg (ref. ^45^), which is suitable for the analyses of interval censored data. The survival dataset consisted of 297 embryos, representative of the three genotypes, sacrificed at age: 3, 4, 5, 7 and 13 days (Supplementary Table 2). Dead embryos constituted left-censored observations (1 < death age ≤ dissection age), whereas live embryos provided right-censored observations (death age > dissection age). We started by fitting semi-parametric proportional hazards models (function ic_sp) and relied on AIC scores to select a best model from a candidate set, including different combinations of predictors (sex and genotype). The final best model featured sex by genotype combinations as predictors. *P*-values were calculated via bootstrapping (n = 10000 samples). Model assumptions were checked using the function *diag_covar*. We then fitted a set of parametric survival models (function *ic_par*), yielding 95% confidence intervals, all having as predictor sex*genotype, but differing in their baseline distribution. We assessed the fit of these different specifications of parametric models by comparing AIC scores and plotting model predictions against the corresponding semi-parametric model fit (Extended Data Fig. 2).

### Selection and processing of chicken embryos and tissues for RNA sequencing analysis

Upon completing the genotyping and sexing of G2 embryos, we selected 36 embryos for RNA sequencing. This selection included 18 E2 embryos (9 males and 9 females) (HH stage 12), 9 E3 males (HH stages 18-19), and 9 E5 males (HH stages 24-25). For the E2 cohort, RNA extraction was performed on whole embryos after the removal of extra-embryonic membranes. This cohort included 9 female embryos of various genotypes (3 ZW, 3 Z^KO^W, and 3 pure HL ZW embryos – i.e., female embryos from the original stock – as a control for maternal effects on gene expression), and 9 male embryos (3 Z^KO^Z, 3 Z^KO^Z^KO^, and 3 ZZ genotypes). Given the low expression of miR-2954 in females and their survival, we then focused on gene expression in males. For the E3 and E5 cohorts, we investigated tissue-specific gene expression by dissecting the head, heart, and rest of the body (referred to as the body) from each male embryo under a stereo microscope, with all dissections performed in ice-cold PBS buffer. Each tissue type from each embryo was represented by 3 replicates derived from 3 individuals, respectively. We note that all ZZ are pure HL and all other genotypes (ZW, Z^KO^W, Z^KO^Z, Z^KO^Z^KO^) are G2.

### RNA extraction and sequencing

A total of 72 samples from E2, E3, and E5 embryos were used for the generation of RNA sequencing libraries. We extracted total RNA from whole embryos or dissected tissues using the AllPrep DNA/RNA/miRNA Universal Kit (Qiagen), following the manufacturer’s protocols. The RNA quality was assessed using the Fragment Analyzer system (Agilent), and all RNA quality numbers (RQNs) were equal to 10, indicating a lack of degradation. The RNA-seq libraries were prepared from 400 ng of RNA per sample using the NEBNext Ultra II RNA Library Prep Kit for Illumina sequencing was performed on an Illumina NextSeq 2000 system, using NextSeq 2000 P3 Reagents (100 Cycles), with samples multiplexed in two sets of 36.

Additionally, we generated small RNA libraries using RNA derived from the same E5 male samples (i.e., those that were also used for the generation of RNA sequencing libraries). This included the generation of small RNA libraries for RNA derived from ZZ (2 replicates), Z^KO^Z (3 replicates), and Z^KO^Z^KO^ (3 replicates) for each tissue type (head, body, heart), respectively. These libraries were prepared using the NEBNext Small RNA Library Prep Set for Illumina and were sequenced on an Illumina NextSeq 550 system using NextSeq 500/550 High Output Kit v2.5 (75 Cycles), with samples multiplexed in two sets of 12.

### Estimation of gene expression levels

The chicken reference genome (bGalGal1.mat.broiler.GRCg7b; GCA_016699485.1) and corresponding GTF annotation file were obtained from Ensembl^46^ (release 109). Raw reads from each library were aligned to the reference genome using STAR aligner v. 2.7.2b (ref. ^47^). This alignment process involved generating STAR indices, aligning reads to the reference genome in an annotation-aware manner, and quantifying the number of reads mapped to each gene using the --quantMode GeneCounts option in STAR. The median uniquely mapped reads number across all samples was 34,703,339. The resulting gene count matrices, along with a metadata file containing sample information and the GTF file, were used to create a RangedSummarizedExperiment object. This object was imported into DESeq2 v. 1.24.0 (ref. ^47^) for downstream analysis. Gene expression data were normalized using Variance Stabilizing Transformation (VST) through the vsn package v. 3.52.0 in R (version 4.1) (ref. ^48^) implemented in the DESeq2 package.

Subsequently, Principal Component Analysis (PCA) was conducted as implemented in the DESeq2 package to examine sample relationships and identify potential outliers. The PCA results revealed a clear clustering of samples (including biological replicates) for the respective tissues and ages without outliers, supporting the high quality of the expression data (Extended Data Fig. 3).

Raw short RNA-sequencing data were preprocessed using a custom Bash script. Adapter sequences were trimmed and reads were size-selected using Cutadapt version 4.4. The parameters set a maximum error rate of 0.25, targeted the adapter sequence “AGATCGGAAGAGCACACGTCTGAACTCCAGTCAC” with a minimum overlap of 6 nucleotides, and allowed no indels, while selecting for read lengths between 19 and 26 nucleotides. After trimming and size selection, the reads were aligned to the chicken reference genome using STAR following the ENCODE miRNA-seq pipeline^49^ (www.encodeproject.org/microrna/microrna-seq-encode4/) (May 2017). This alignment process included mapping to the microRNA subset of the chicken GTF gene annotation and quantifying the number of aligned reads in STAR. The median of the number of uniquely mapped reads across all samples was 432,069.

### MicroRNA target prediction in chicken and zebra finch

To identify potential targets of miR-2954, we utilized TargetScan^24^, which detects 6mer, 7mer-1a, 7mer-m8, and 8mer-1a target sites in the 3’ untranslated regions (3’ UTRs) of mRNA transcripts, aligning them with the microRNA seed sequence. We obtained 3’ UTR sequences for all splice variants of genes within both the chicken (bGalGal1.mat.broiler.GRCg7b) and zebra finch (bTaeGut1_v1.p) genomes using BioMart (ref. ^50^). Subsequent identification of target sites was performed using TargetScan version 7 for each species. A gene was categorized as a predicted target if it contained any of these target site types within its UTRs. Additionally, we quantified the total number of target sites for every predicted target gene for the chicken (Supplementary Table 3).

### Differential gene expression analysis

The 3’ UTR is specific to protein-coding genes, and microRNA targets are predicted based on the presence of target sites within their 3’ UTRs. Consequently, we limited the DESeq2 dataset to protein-coding genes (as identified in the GTF annotation). Differential expression analysis was conducted using DESeq2. Differentially expressed genes were identified using a threshold of less than 0.05 for *P*-values, adjusted according to the Benjamini-Hochberg (BH) method^51^. The effect of genotype on gene expression in E2 whole embryos was independently analyzed in male and female embryos (model: gene expression ∼ genotype). For each tissue (head, heart, body), gene expression analysis was performed collectively across ages, employing a model that included both genotype and embryonic age as variables (model: gene expression ∼ genotype + embryonic age). Log fold changes and differentially expressed genes were determined for each genotype contrast (Supplementary Table 4).

### Differential microRNA expression analysis

Differential expression analysis was conducted using DESeq2. Differentially expressed microRNAs were identified using a threshold of less than 0.05 for *P*-values, adjusted according to the Benjamini-Hochberg (BH) method^51^. The effect of genotype on microRNA expression in E5 head, heart, and body was independently analyzed in each tissue (model: gene expression ∼ genotype) (Supplementary Table 5).

### Comparison of pure HL females with ZW G2s

Although all chickens used in the gene expression analysis were of the HL breed, the G2 animals, comprising genotypes ZW, Z^KO^W, Z^KO^Z, and Z^KO^Z^KO^, originated from different parents than the ZZ genotype, which was derived from the pure HL breed (i.e., the original stock). To ensure the rigor of all expression comparisons, we aimed to confirm that the G2 ZW and the pure HL ZW had similar gene expression profiles (N.B.: ZZ embryos cannot be derived from the G2 – hemizygous/heterozygous KO – parents), thereby eliminating potential confounding factors, such as maternal effects on gene expression. Accordingly, we conducted different expression analyses between pure HL ZW and G2 ZW chickens and compared the fold changes across different gene categories. This analysis confirmed that gene expression patterns are statistically indistinguishable between HL and G2 and therefore do not confound our results (Extended Data Fig. 6).

### Identifying ohnologs

The list of chicken ohnologs was retrieved from the OHNOLOGS v2 database^52^, available at http://ohnologs.curie.fr/ (“relaxed” dataset). These ohnologs were identified using gene IDs from the galGal4 assembly (Ensembl release 80), which is incompatible with the gene IDs of the chicken genome assembly employed in our study (GRCG7b). To resolve this, we retrieved the unspliced DNA sequences of these ohnolog gene IDs from the GRCg6a assembly (Ensembl release 106) via BioMart. Subsequently, these sequences were aligned to the unspliced DNA sequences of protein-coding genes from the GRCG7b assembly using BLASTn (BLAST+ 2.4) (ref. ^53^), with the settings -perc_identity 95 and -evalue 0.001. We sorted the results by bit scores to identify the best hits between the two gene sets. Cross-referencing protein names for matched gene IDs confirmed a high accuracy (88.6% exact matches) of this ID conversion method (Supplementary Table 3).

### Dosage sensitivity scores

Dosage sensitivity scores for human genes, including haploinsufficiency (pHaplo) and triplosensitivity (pTriplo), were sourced from a previous study^30^. These scores were then assigned to chicken genes based on their 1:1 orthology relationship (retrieved using BioMart) (Supplementary Table 3).

### Assessment of time- and tissue-specificity

To evaluate the time and tissue-specificity of chicken genes, we calculated time and tissue-specificity indexes based on the Tau metric^54^ using a developmental time-series RNA-seq dataset^32^ (Supplementary Table 6). As in previous studies^32^, for the tissue-specificity index, the Tau metric was applied to the maximum expression of the gene observed during development in each organ, whereas for the time-specificity index, the Tau metric was applied to the expression of the gene at different time points instead of organs. In both cases, indexes range from 0 (indicating broad expression) to 1 (indicating restricted expression).

### Female to male expression level ratios

Gene expression ratios between the sexes were analyzed using a published RNA-seq time-series dataset^32^. We obtained raw read (fastq) files for various chicken organs (brain, cerebellum, heart, kidney, liver) across different embryonic stages (E10, E12, E14, E17) and post-hatch periods (P0, P7, P35, P70, P155). Reads were aligned to the bGalGal1.mat.broiler.GRCg7b reference genome, with read counts generated as detailed in Section 7. We then calculated FPKM values for each gene using the fpkm function in DESeq2 and determined median expression values for all embryonic and all post-hatch samples (Supplementary Table 6). An FPKM threshold greater than 1, based on the median for each group, was applied to filter out non-expressed and lowly expressed genes in both sexes.

### Assessment of Z to proto-Z expression levels

For this analysis, RNA-seq data (log2 transformed RPKM values from ref. ^32^) from brain, cerebellum, heart, kidney and liver from adult male and female chicken (P155) and the corresponding stage in mice (P63) was used. Akin to previous studies^5,6,9^, ancestral expression levels of Z-linked genes (proto-Z genes) were estimated by calculating the median expression levels of the corresponding expressed autosomal 1∶1 orthologs in an outgroup species with non-ZW sex chromosomes (in this case mouse). In a similar way, ancestral expression levels of autosomal genes (proto-autosomal genes) were estimated by calculating the median expression levels of corresponding 1∶1 orthologs that are autosomal in the same outgroup species with non-ZW sex chromosomes.

To obtain the current-Z to proto-Z expression ratios we first normalized the current expression levels of Z-linked genes by the median current expression level of all 1∶1 orthologous genes that are autosomal in the outgroup species. We then normalized the ancestral expression levels of each proto-Z–linked gene (computed as described above) by the median ancestral expression level of all proto-autosomes in the outgroup species. We then computed the ratio of these two values for each gene, resulting in the current-Z to proto-Z ratios.

Finally, we compared the current-Z to proto-Z ratios for Z-linked miR-2954 targets and Z-linked miR-2954 non-targets. As Z-linked targets, we used the experimental miR-2954 targets and as non-targets we used Z-linked genes that are neither experimental miR-2954 targets nor predicted miR-2954 targets. In both cases, we made sure that autosomal miR-2954 targets were excluded when normalizing the expression of current-Z and proto-Z genes by current-autosomal and proto-autosomal genes. Statistically significant deviations of the medians of these ratios from key reference values (e.g., 0.5 [log2 ratio of −1]; 1 [log2 ratio of 0]; and 2 [log2 ratio of 1]) were assessed using one-sample Wilcoxon signed rank tests. *P*-values were corrected for multiple testing using the Bonferroni procedure^55^, with adjusted *P* < 0.05 indicating significance.

### Location of genes along the Z chromosome

To visualize the location of target genes on the Z chromosome, we counted the number of protein coding genes in windows of 0.5 Mb based on gene annotations of Ensembl^46^ (version 111). To indicate the location of the male hypermethylated regions, we used the regions defined by Sun et al. (ref. ^35^). We lifted these regions from Galgal5.0 to the bGalGal1.mat.broiler.GRCg7b genome assembly by extracting flanking sequences from and aligning them to the new genome with BLAT.

### Sequence conservation

The sequence of the miR-2954 locus was retrieved from NCBI and blasted against the reference genomes of the target species (Extended Data Fig. 5) using BLASTn (ref. ^53^).

## SUPPLEMENTARY INFORMATION

**Supplementary Table 1.** Oligonucleotide sequences used in this study, including primers for sexing and genotyping, as well as single guide RNA (sgRNA) sequences and the single-stranded DNA oligonucleotides (ssODNs) repair template.

**Supplementary Table 2**: Dataset on the viability screening of all embryos utilized in this study, detailing the embryonic age at dissection, sex, genotype, and vitality status. Genotypes are defined as ’WT’ for wild-type, ’HZ’ for heterozygous, ’KO’ for homozygous in males, and hemizygous in females.

**Supplementary Table 3**. Dataset with the following information: each gene’s unique identifier (gene_id); chromosome names where the genes are located (seqnames); common names or symbols for each gene (gene_name); classification of genes by genomic features and biological roles (gene_biotype); counts of microRNA binding sites predicted by TargetScan, including site_6mer, site_7mer_1a, site_7mer_m8, and site_8mer_1a, which vary in complementarity; stable identification numbers for human genes that are 1:1 orthologous (human_gene_stable_ID); official gene symbols for these human orthologs (human_gene_name); scores indicating the probability of haploinsufficiency or triploinsufficiency (pHaplo, pTriplo); gene identifiers according to the chicken reference genome version GalGal6a (gal6_gene_id); an indicator of whether the gene is among chicken ohnologs (if_ohno); whether the gene is among experimentally validated targets of miR-2954, as identified in this study (expr_targets); identifiers for 1:1 orthologous genes found in the Taeniopygia guttata (zebra finch) genome (zebra_finch_gene_id); and an indicator of whether the gene is a predicted target of miR-2954 in the zebra finch (zebra_finch_target).

**Supplementary Table 4:** Dataset presenting the DESeq2 analysis results for various contrasts involving E2 whole embryos and embryonic day 3-5 (E3-5) tissues from the head, heart, and body. It also includes annotations indicating whether the genes are located on the Z chromosome or autosomes (’chr_z_or_auto’) and whether they are predicted targets of miR-2954 (’mir_target_group’).

**Supplementary Table 5:** Dataset presenting the standard output from a DESeq2 analysis comparing the expression of microRNAs in male embryonic day 5 tissues from the head, heart, and body when homozygous individuals are contrasted with wild-types.

**Supplementary Table 6:** Dataset of FPKM (Fragments Per Kilobase of transcript per Million mapped reads) values, profiling gene expression across various tissues including the cerebellum, brain, heart, liver, and kidney. The developmental stages covered are embryonic days 10, 12, 14, and 17, as well as postnatal days 0, 7, 35, and 155. For each combination of tissue and developmental stage, the dataset provides information from two biological replicates.

**Supplementary Data 1:** Photographs of embryos used in gene expression analysis and/or viability screening, captured using a stereo microscope.

---

**Extended Data Fig. 1.**
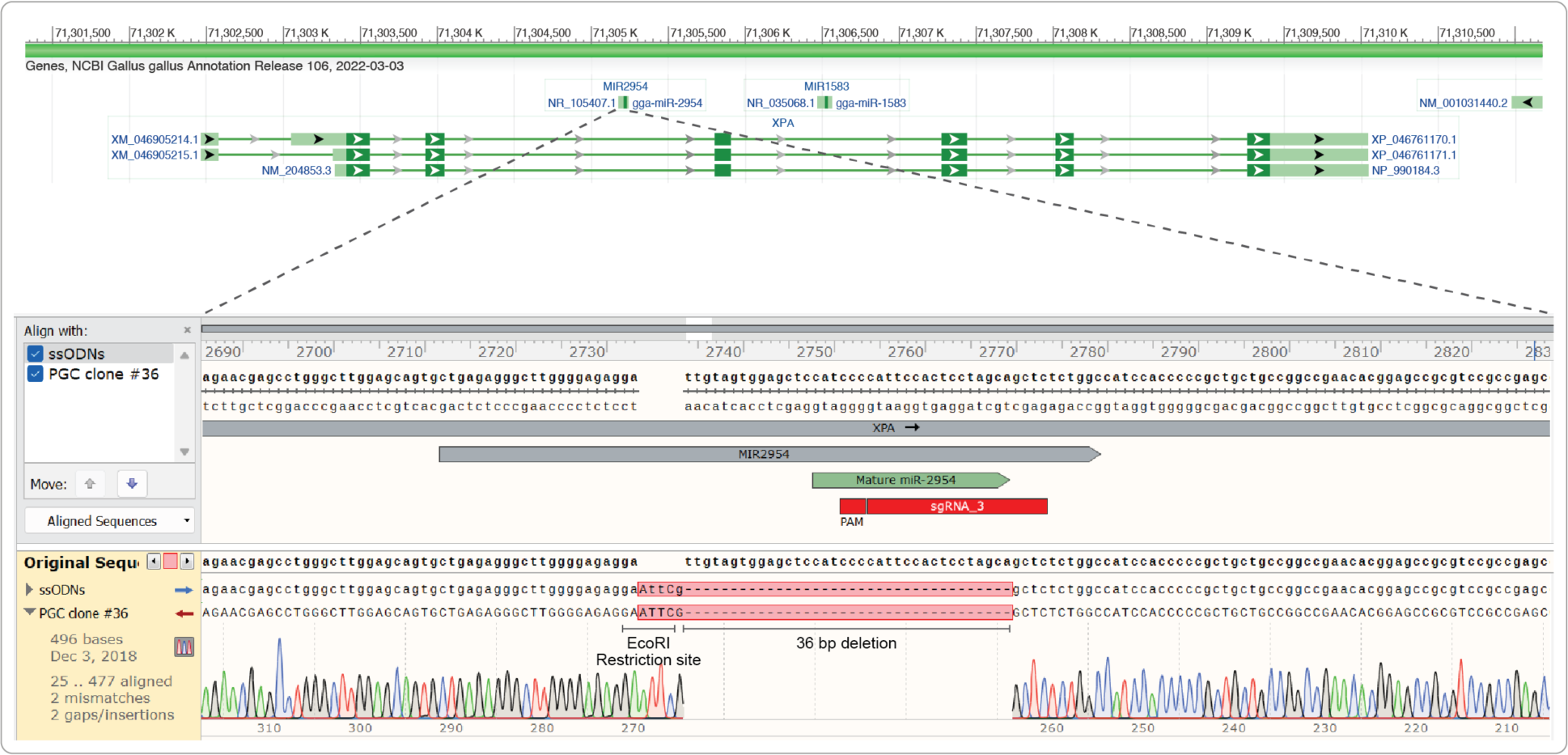
Overview of the knockout of the miR-2954 locus. Top: overview of the *XPA* host gene, with the location of the miR-2954 locus indicated in the second or third (depending on the isoform) intron of this gene. Bottom/middle graphs: alignments/overview of the genomic reference sequence around miR-2954 locus, the induced deletion, and the single-stranded DNA oligonucleotide (ssODN) repair template utilized in this study to exploit the homology-directed repair (HDR) pathway (Methods). Sequence track with the locations of pre- and mature miR-2954 sequences. The ssODN repair template aligns with the post-editing PGC clone sequence, as confirmed by Sanger sequencing (clone #36, used for generating KO chickens), highlighting a 36 bp deletion adjacent to an EcoRI restriction site. The accompanying chromatogram verifies the deletion, illustrating the consistency between the edited and expected sequences.

**Extended Data Fig 2.**
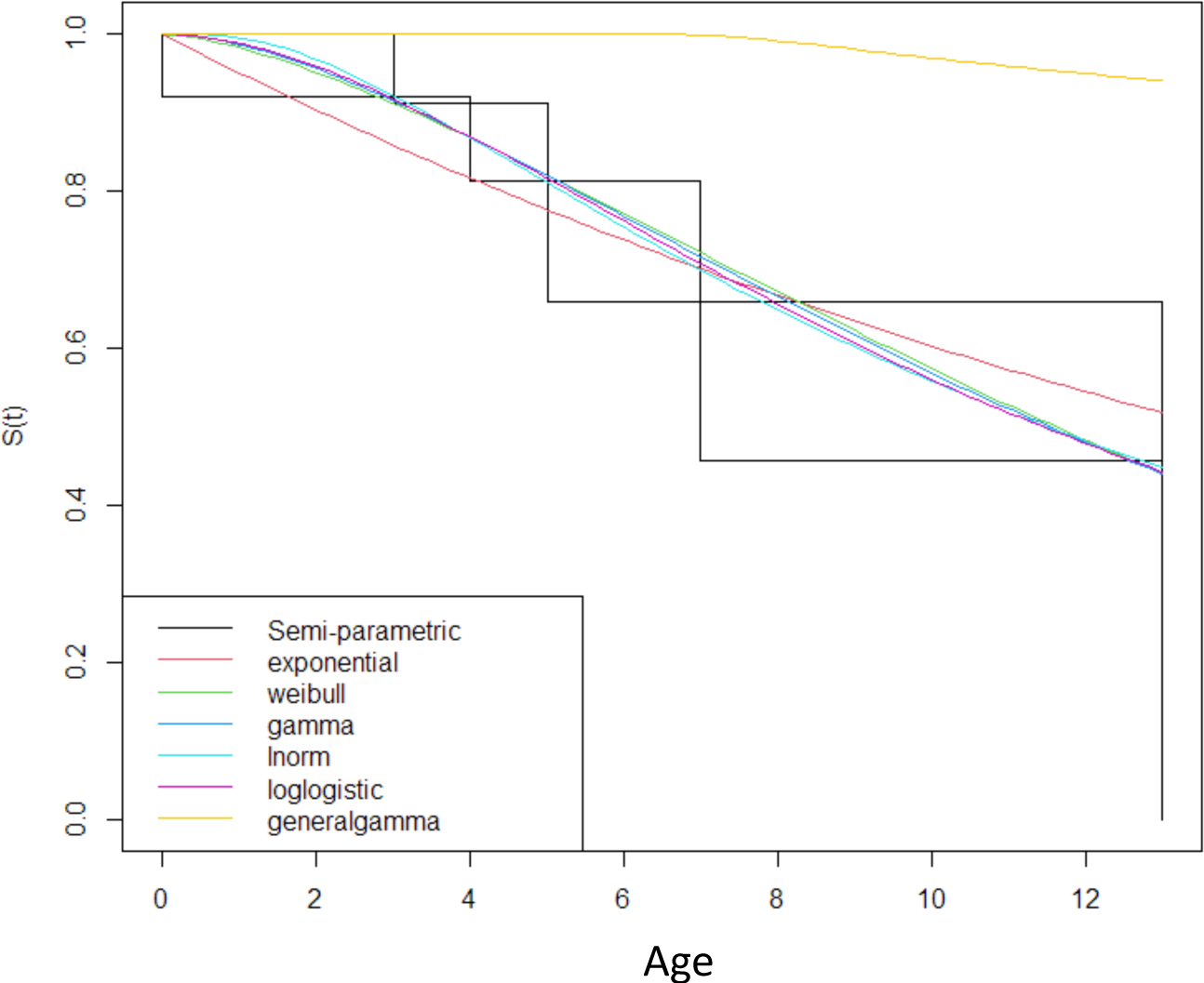
Assessment of the fit of different specifications of parametric survival models. Parametric baseline fits for different distributions (represented by different colors) compared to semi-parametric model fit (in black). Most distributions fitted similarly well, with the exception of exponential and general gamma functions. The final parametric model, selected based on AIC scores, featured a log-normal (“lnorm”) distribution.

**Extended Data Fig 3.**
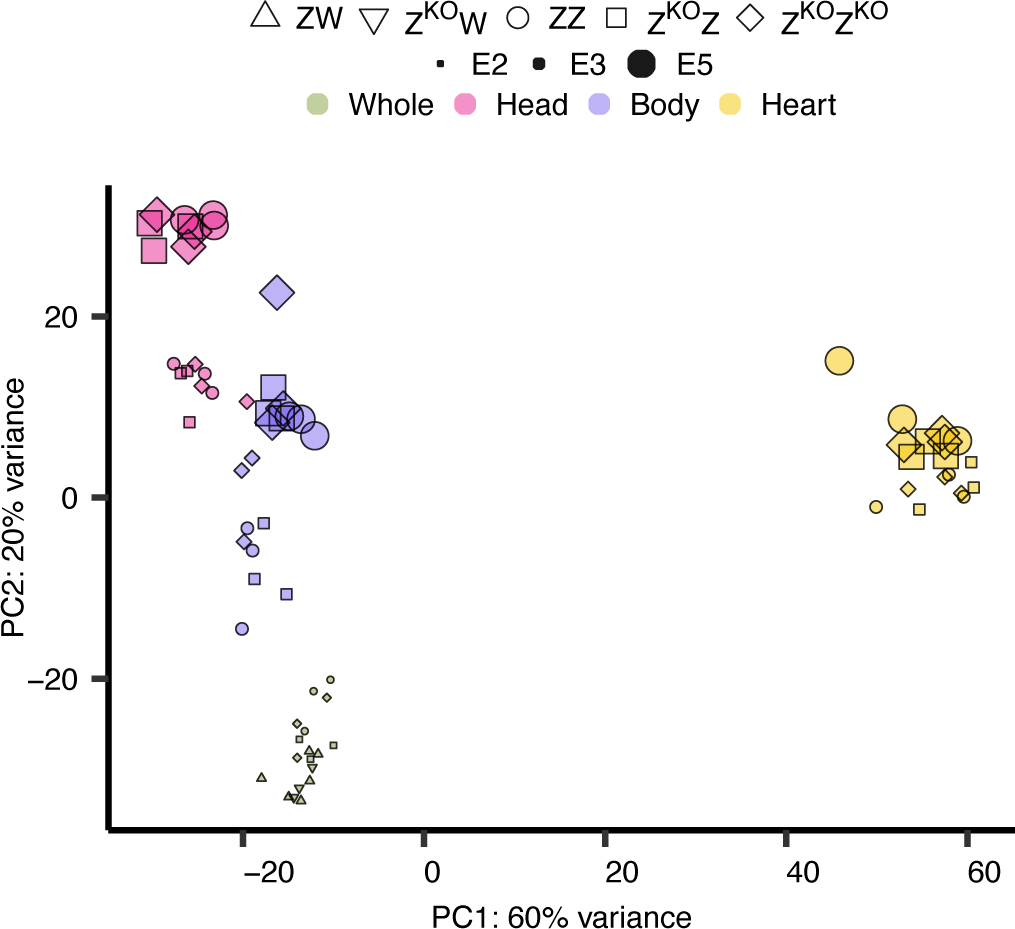
Principal component analysis (PCA) across of mRNA expression levels across samples. The proportion of the variance explained by the two first principal components (PC1 and PC2) is indicated.

**Extended Data Fig 4.**
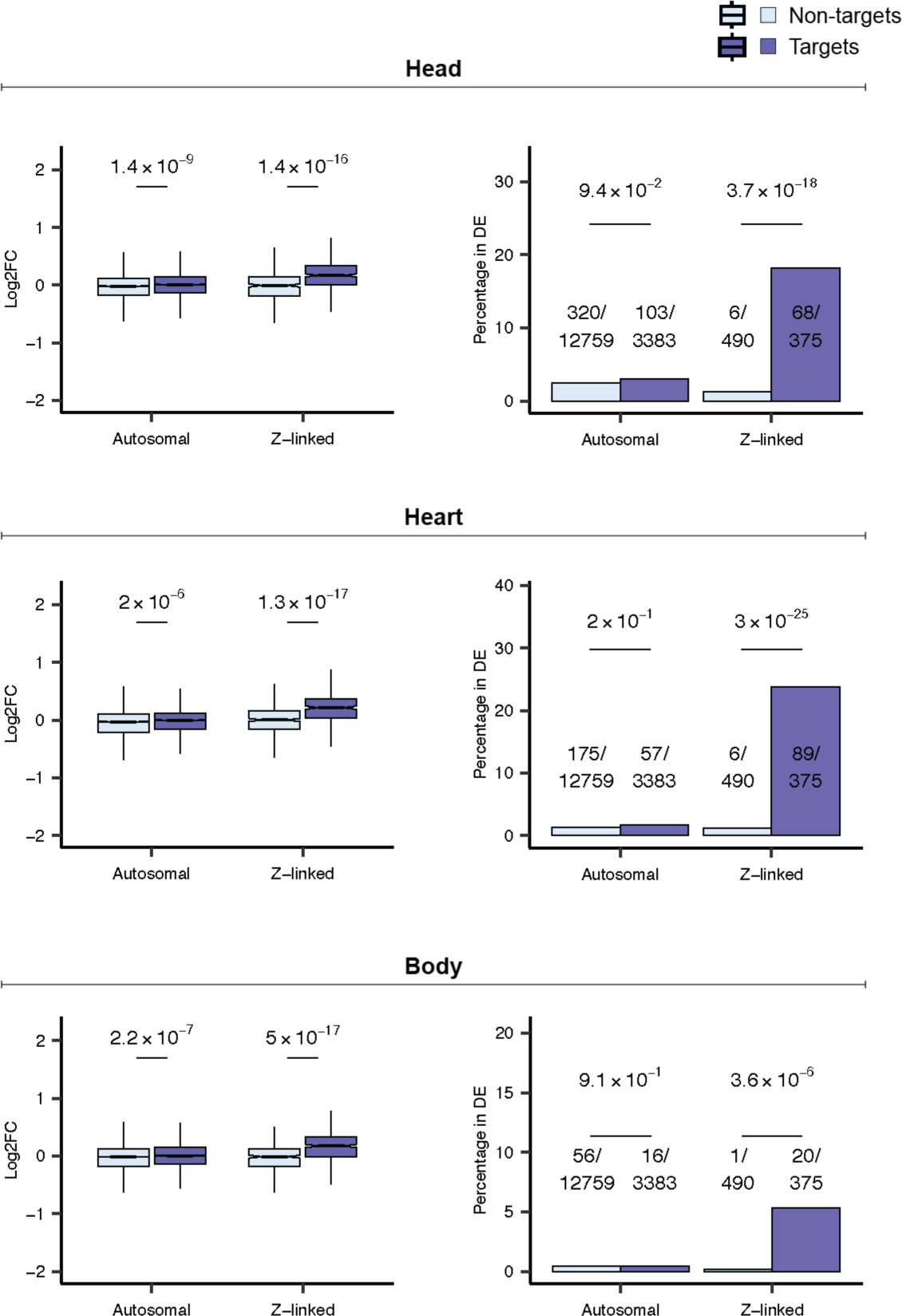
Impact of miR-2954 knockout on gene expression in heterozygous KO males. Left column: log_2_-fold changes (Log2FC) in gene expression between Z^KO^Z and ZZ genotypes for both autosomal and Z-linked protein-coding genes; *P*-values from a two-sided Wilcoxon rank-sum test are indicated above. Log2FC estimates are based on transcriptomes of E3 and E5 embryos for the head, heart, and rest of the body, respectively. Right column: proportions of autosomal and Z-linked target and non-target genes among the differentially expressed (DE) genes (Benjamini-Hochberg adjusted *P* < 0.05) when contrasting Z^KO^Z and ZZ genotypes. The distribution of predicted targets and non-targets of miR-2954 is compared using a chi-squared test for each group.

**Extended Data Fig 5.**
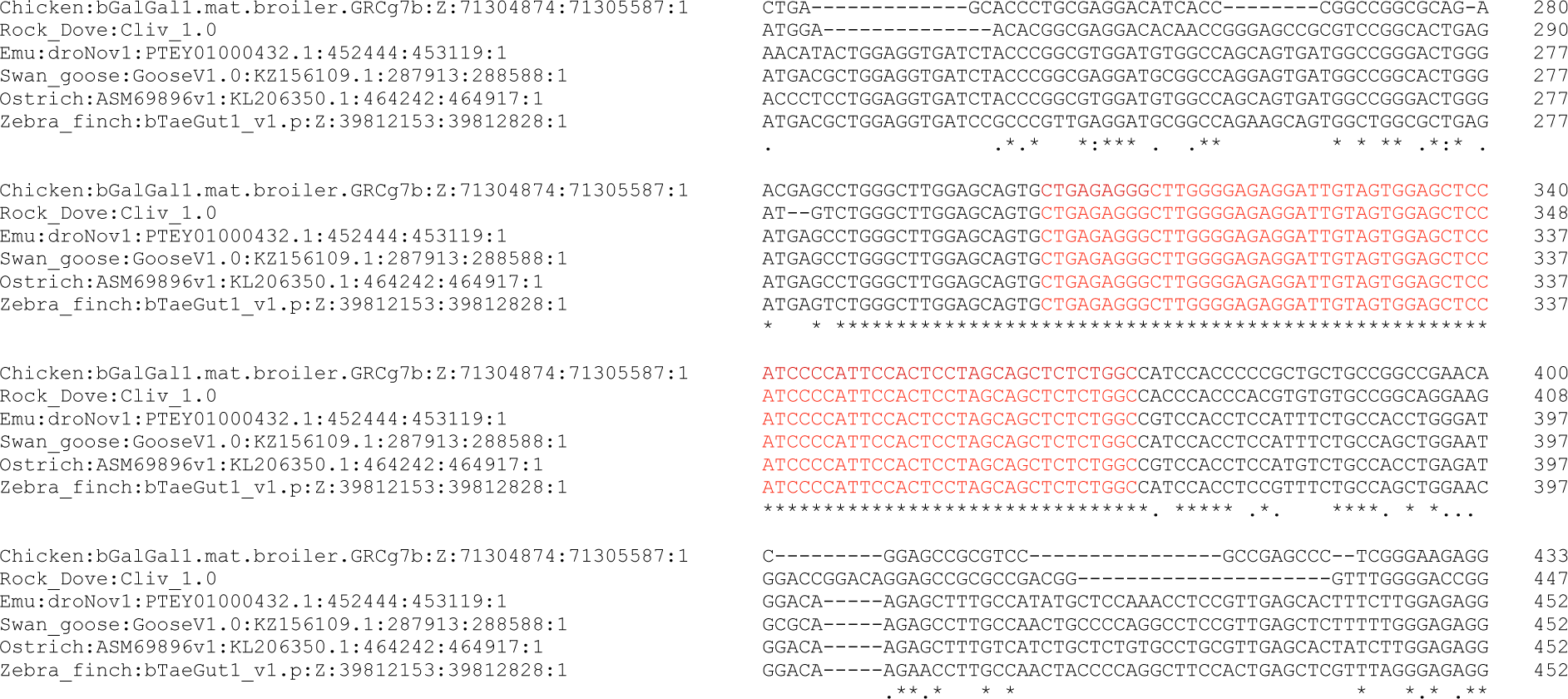
Alignment of the miR-2954 locus sequence across representative bird species. CLUSTAL O (1.2.4) multiple sequence alignment, displaying a segment of the second/third intron of the *XPA* gene, which includes the miR-2954 genetic sequences from representative bird species. Dashes signify sequence gaps introduced to optimize the alignment. The consensus line below the alignment denotes residues with identical nucleotides across species (’*’) and those where nucleotides are partly shared between species (’:’, ’.’).

**Extended Data Fig 6.**
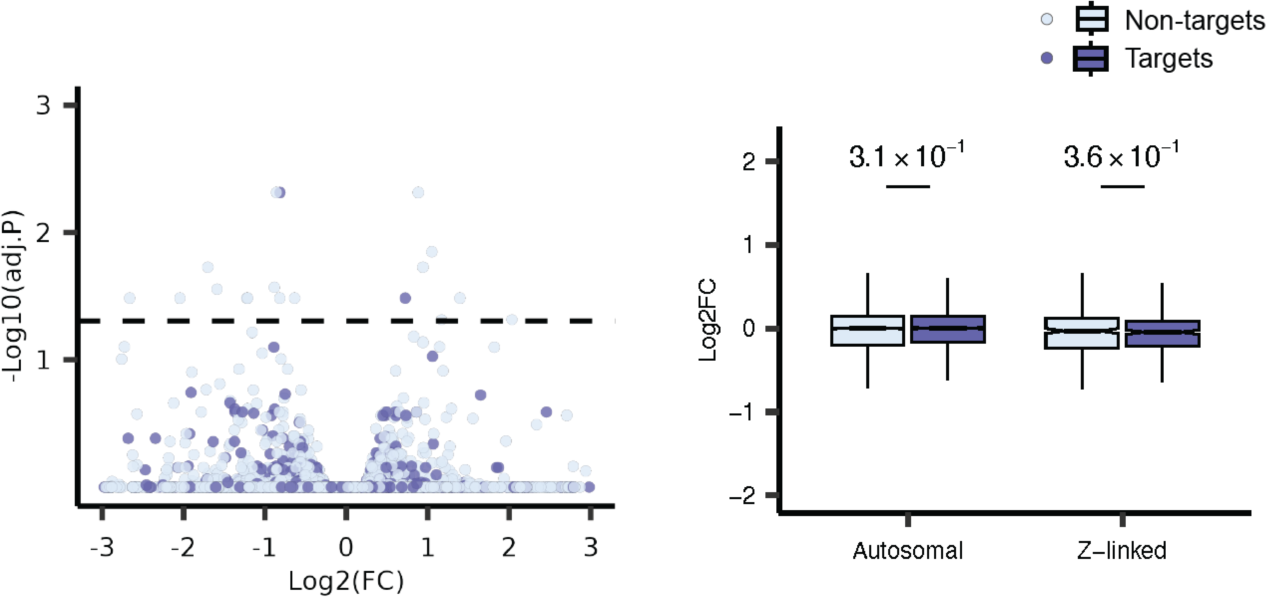
Comparison of “pure” HL females (from original stock) with ZW females from G2. Left panel: The log2FC and -log10 of Benjamini-Hochberg adjusted p-values for predicted miR-2954 target and non-target genes, contrasting female pure Hy-Line and generation 2 (G2) wild-type (ZW) females. Right panel: comparison of the log2 fold change (Log2FC) in gene expression between pure Hy-Line and generation 2 (G2) wild-type (ZW) females within both autosomal and Z-linked categories for protein-coding genes, with p-values from a two-sided Wilcoxon rank-sum test indicated above the boxplots. Log2FC estimates are based on the transcriptome of embryonic day 2 whole embryos.

