## Supplementary figures and images for "A male-essential microRNA is key for avian sex chromosome dosage compensation"

### Supplementary Data 1

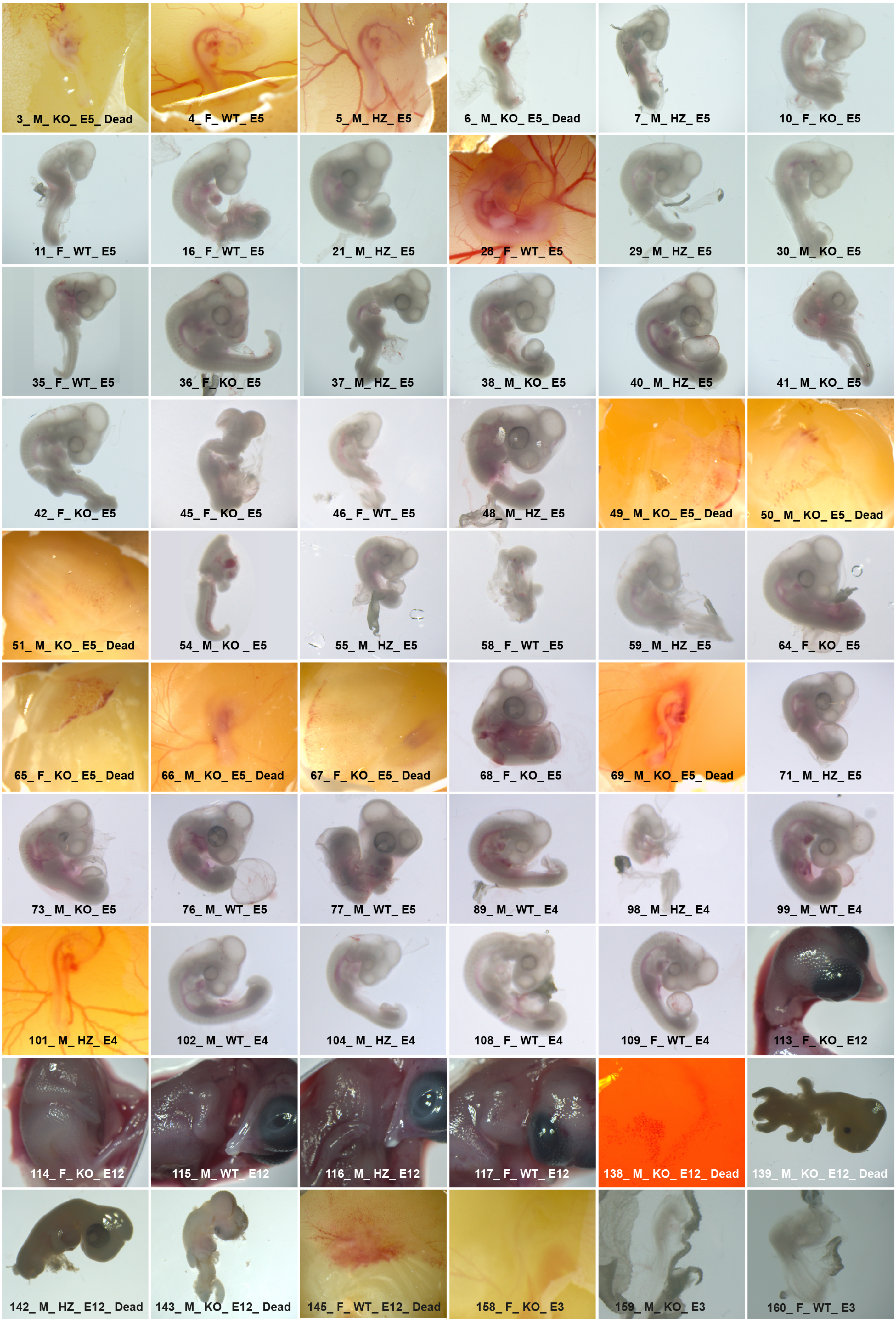

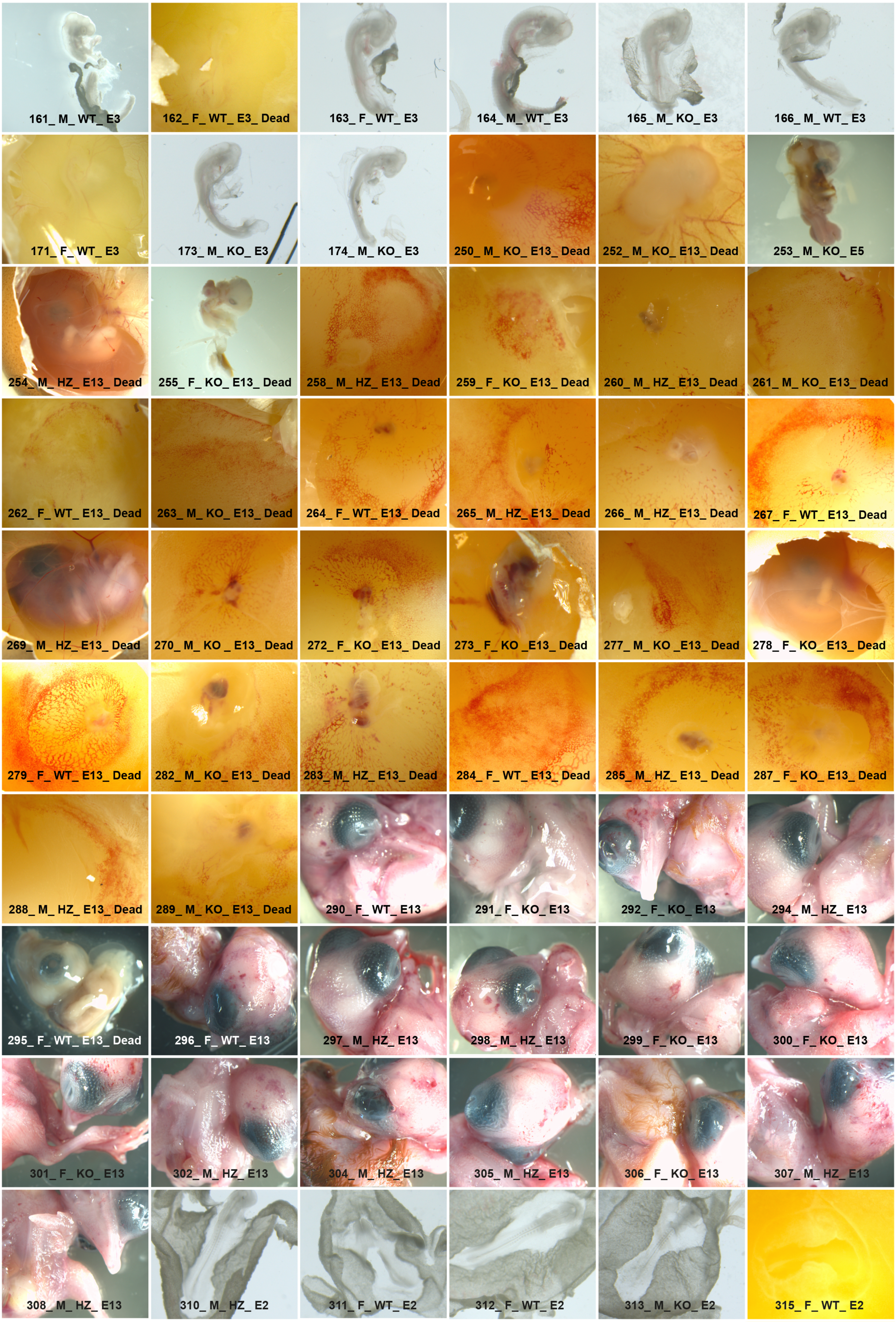

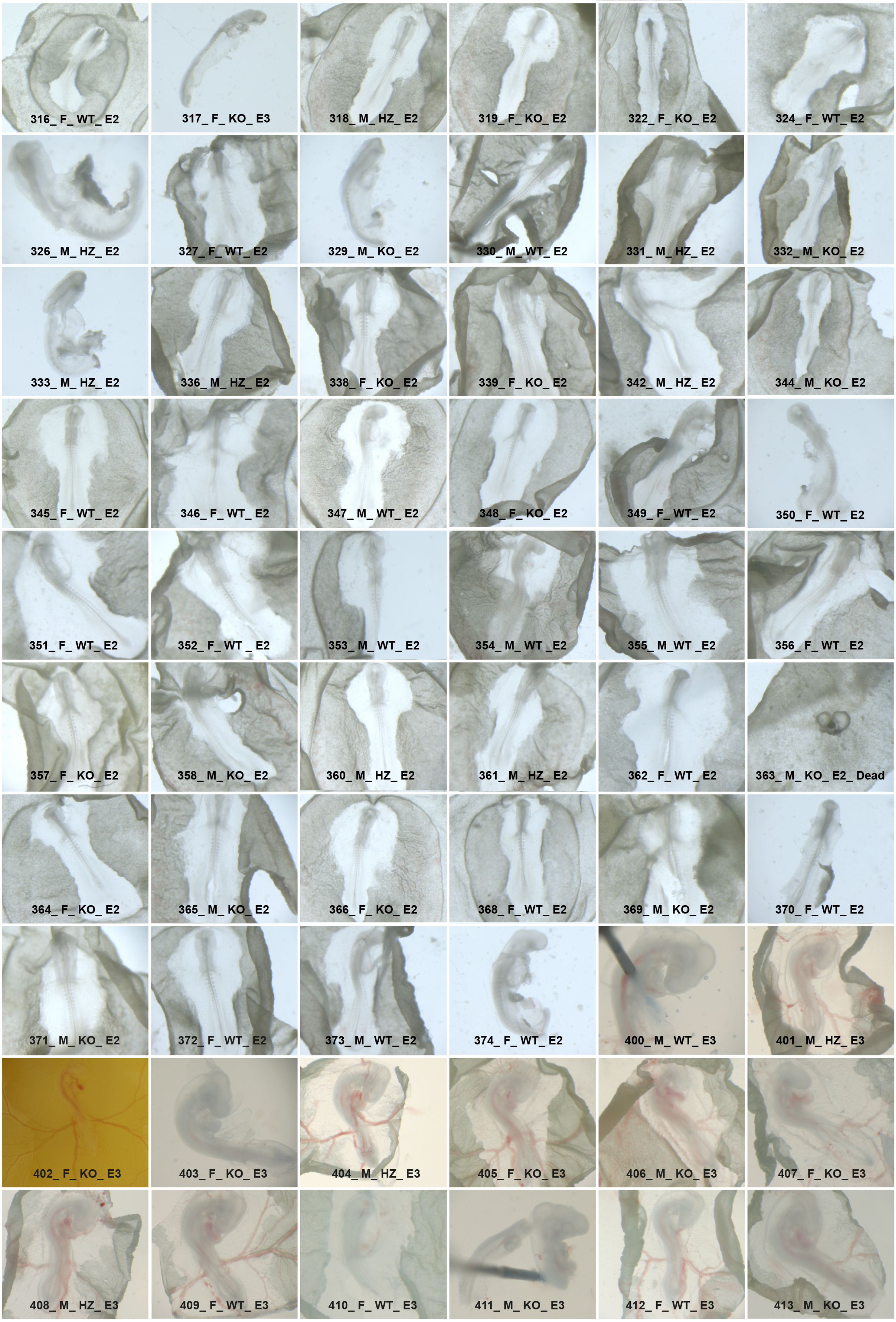

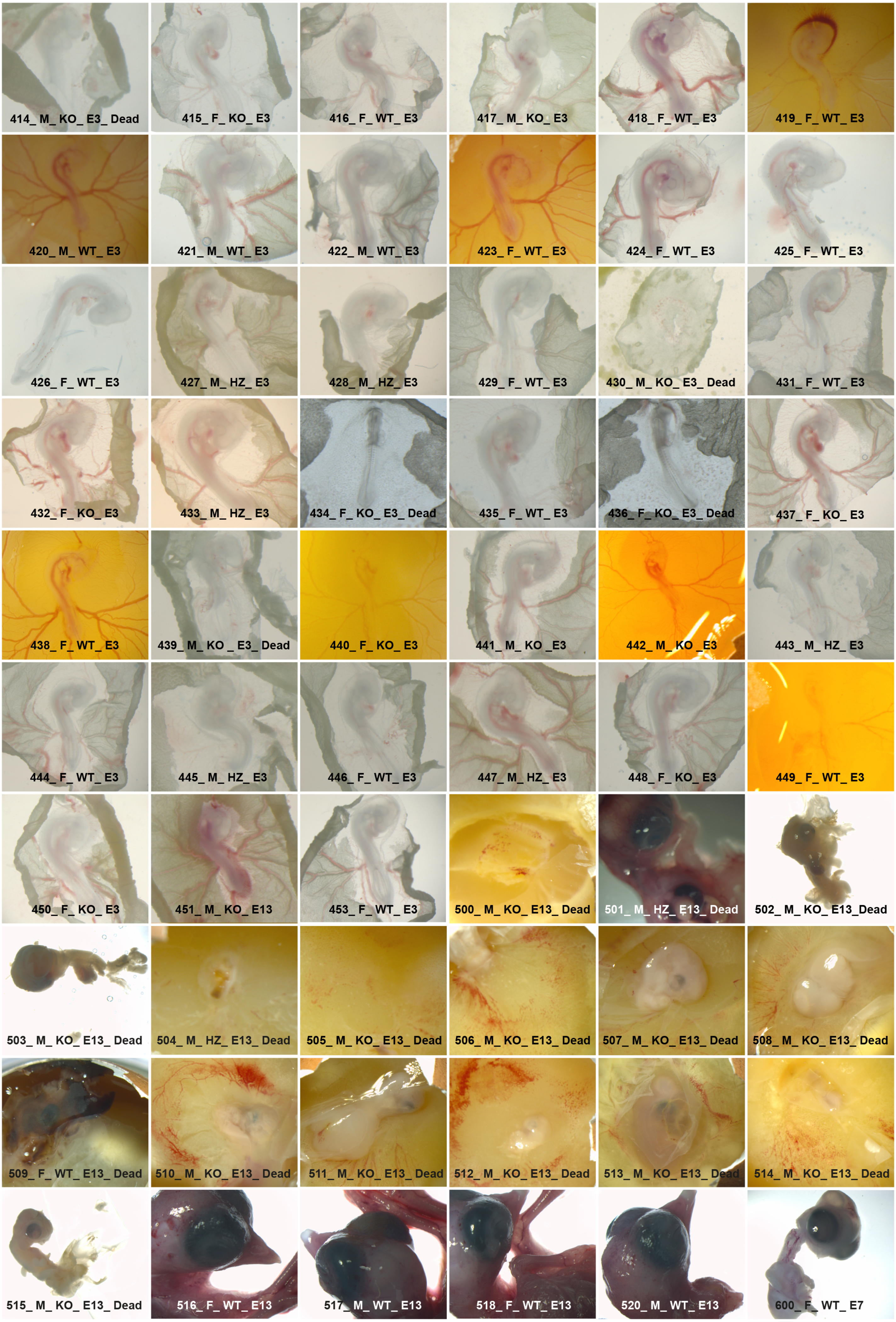

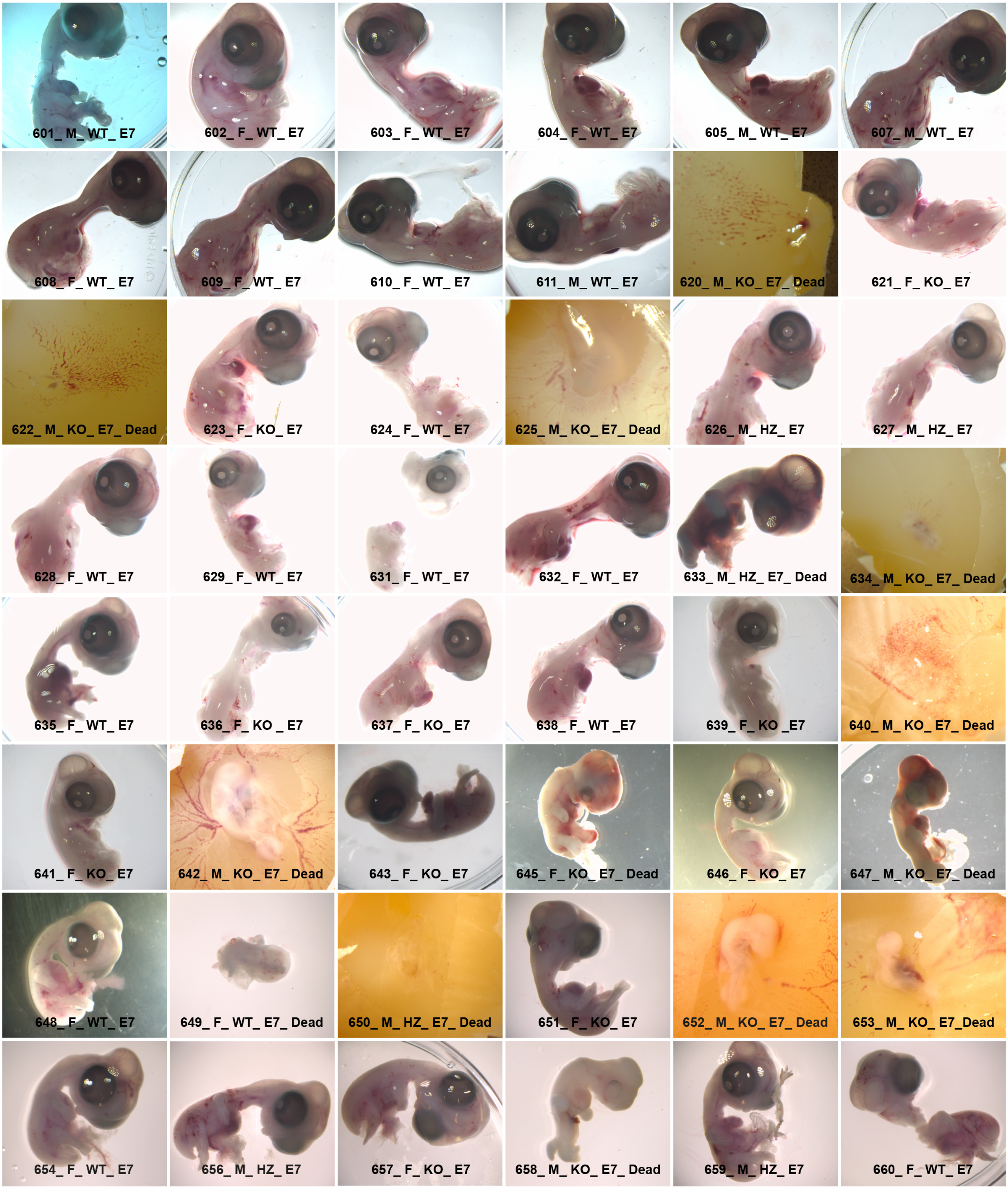

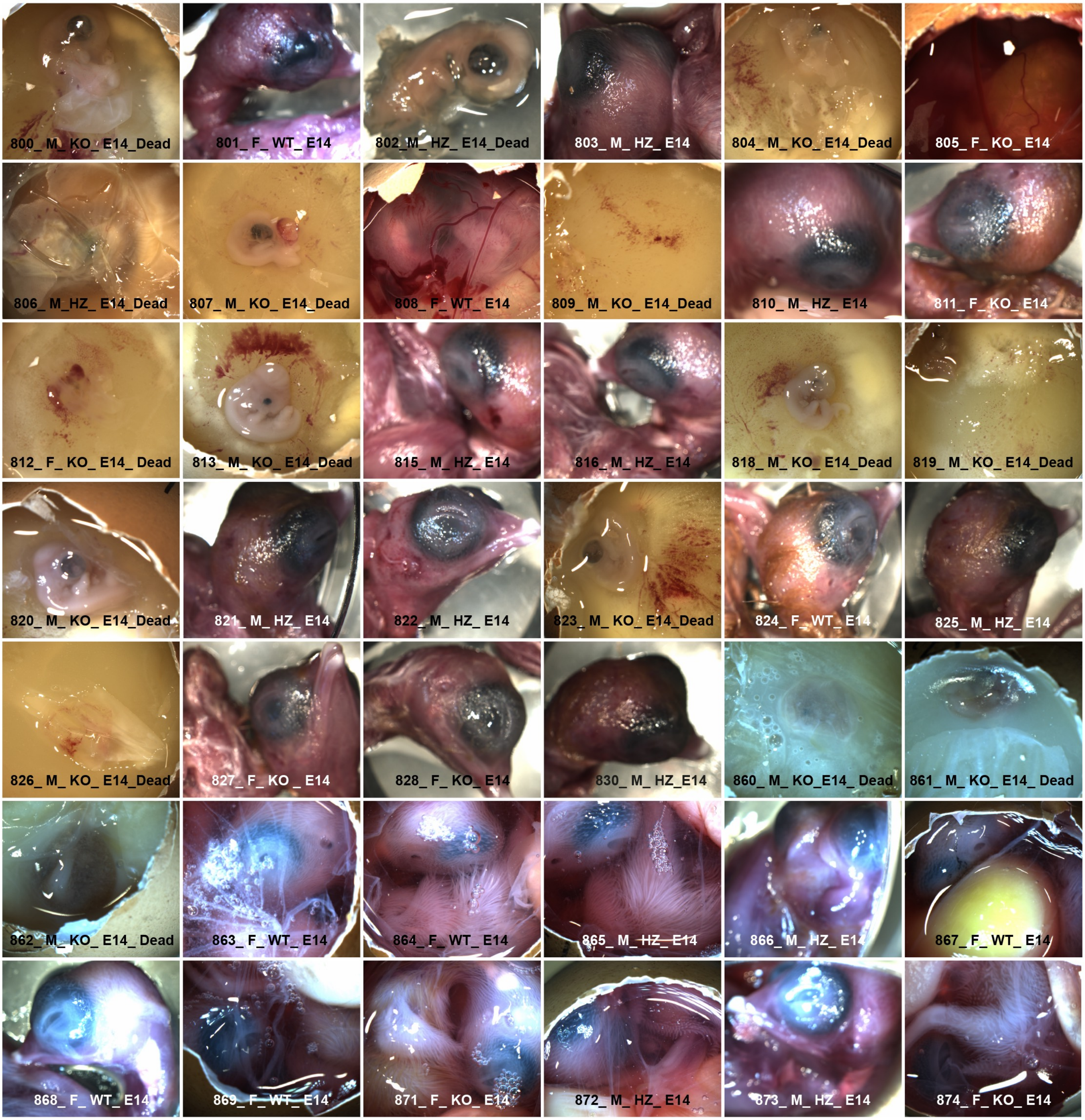
