## Supplementary Table 1 for "A male-essential microRNA is key for avian sex chromosome dosage compensation"

| Primer name | Sequence (5’ to 3’) | Product (bp) | Application |
| --- | --- | --- | --- |
| 12S (F) (Control) | CTATAATCGATAATCCACGATTCA | 131 | Sexing control |
| 12S (R) (Control) | CTTGACCTGTCTTATTAGCGAGG |  |  |
| Sex_1 (F) (W-Specific) | GAGATCACGAACTCAACCAG | 211 | Sexing W-specific |
| Sex_1 (R) (W-Specific) | CCAGACCTAATACGGTTTTACAG |  |  |
| XPA-INT2-F4 | AATGAGAGCTGTGTAAGGCGT | 550 | Genotyping |
| XPA-INT2-R4 | GCTACGAAATGAAGCACGACG |  |  |
| sgRNA_3_Top | caccGAGCTGCTAGGAGTGGAATG | - | Short guide RNA (sgRNA targeting miR-2954 |
| sgRNA_3_Bottom | aaacCATTCCACTCCTAGCAGCTC |  |  |
| ssODNs_template | gcgcagaacgagcctgggcttggagcagtgctgagagggcttggggagaggaAttCggctctctggccatccacccccgctgctgccggccgaacacggagccgcgtccgccgagcc | - | Single-stranded DNA oligonucleotides (ssODNs) repair template |

Supplementary Table 1. Oligonucleotides used in this study.
